# Multiple bursts of speciation in Madagascar’s endangered lemurs

**DOI:** 10.1101/2023.04.26.537867

**Authors:** Kathryn M. Everson, Luca Pozzi, Meredith A. Barrett, Mary E. Blair, Mariah E. Donohue, Peter M. Kappeler, Andrew C. Kitchener, Alan R. Lemmon, Emily Moriarty Lemmon, Carlos J. Pavón-Vázquez, Ute Radespiel, Blanchard Randrianambinina, Rodin M. Rasoloarison, Solofonirina Rasoloharijaona, Christian Roos, Jordi Salmona, Anne D. Yoder, Rosana Zenil-Ferguson, Dietmar Zinner, David W. Weisrock

## Abstract

Lemurs are often cited as an example of adaptive radiation. Since colonizing Madagascar, more than 100 extant lemur species have evolved to fill a variety of ecological niches on the island. However, recent work suggests that lemurs do not exhibit one of the hallmarks of other adaptive radiations: explosive speciation rates that decline over time. Thus, characterizing the tempo and mode of evolution in lemurs can help us understand alternative ways that hyperdiverse clades arise over time, which might differ from traditional models. We explore the evolution of lemurs using a phylogenomic dataset with broad taxonomic sampling that includes the lemurs’ sister group, the lorisiforms of Asia and continental Africa. Our analyses reveal multiple bursts of diversification (without subsequent declines) that explain much of today’s lemur diversity. We also find higher rates of speciation in Madagascar’s lemurs compared to lorisiforms, and we demonstrate that the lemur clades with exceptionally high diversification rates have higher rates of genomic introgression. This suggests that hybridization in these primates is not an evolutionary dead-end, but a driving force for diversification. Considering the conservation crisis affecting strepsirrhine primates, with approximately 95% of species being threatened with extinction, this phylogenomic study offers a new perspective for explaining Madagascar’s exceptional primate diversity and reveals patterns of speciation, extinction, and gene flow that will help inform future conservation decisions.

## Main Text

The lemurs of Madagascar (Strepsirrhini: Lemuriformes and Chiromyiformes^1^) are a fascinating case study in evolutionary biology. They are exceptionally diverse—representing more than 15% of all living primate species—yet all members of the clade live on an island representing < 1% of Earth’s land area^2^. After colonizing Madagascar, lemurs evolved to fill a wide range of ecological niches, from the smallest primate species in the world—the arboreal mouse lemurs (*Microcebus*)—to recently extinct terrestrial species as large as female gorillas (*Archaeoindris*). Given their exceptional phenotypic and ecological diversity, lemurs are often highlighted as an example of adaptive radiation^3^ along with other classic examples like Darwin’s finches from the Galápagos Islands^4^ and cichlids from Lake Victoria^5^. However, a recent study^6^ found that lemurs did not follow an expected pattern of adaptive radiation, i.e., they did not experience rapid or explosive speciation that decreased over time as niches became filled^7,8^. With this new understanding of the overall rates of lemur diversification, the stage is set to further unravel evolutionary tempo in the accumulation of their exceptional diversity. Access to genomic data provides the opportunity to refine estimates of lemur phylogeny and branch lengths, to test more detailed models of diversification, and to ask whether previously unexplored evolutionary factors have shaped lemur diversity.

To fully understand the evolutionary dynamics of lemurs, we must properly contextualize their diversification alongside their often-neglected sister group, Lorisiformes. Lemurs and lorisiforms (collectively known as the “wet-nosed primates,” suborder Strepsirrhini) together form an excellent comparative system for understanding how evolutionary dynamics in different geographical regions can produce drastically different levels of species diversity. The lorisiform primates, which occur in Asia and continental Africa, include galagos, pottos, angwantibos, and lorises, all of which are nocturnal and elusive. While they exhibit several interesting morphological adaptations—e.g., they include the only venomous primates (*Nycticebus* and *Xanthonycticebus*)—the lorisiforms are less diverse than lemurs overall, both phenotypically and in terms of species diversity^6,9^. As a result, they have been comparatively neglected in scientific literature^10^. Here, we use a phylogenomic dataset to reconstruct the evolutionary history of Strepsirrhini, providing a framework for evaluating if lemurs diversified according to the classic adaptive radiation model and whether their rates of diversification differed from those of lorisiforms. Given abiding uncertainty about phylogenetic relationships within these groups (discussed in the following section), we also consider the possibility that introgressive hybridization has introduced conflicting genealogical histories across the genome. Hybridization has been historically conceptualized as a homogenizing force in evolutionary biology that counteracts divergence^11^, but a recent systematic review of adaptive radiations showed that gene flow often provides fuel for diversification as well^12^. To address this idea, we additionally test for a relationship between introgression and the rate of diversification in strepsirrhines, providing insights into a continuing question in evolutionary biology, i.e., whether hybridization impedes or promotes the formation of new species ^13–15^.

## Results and Discussion

### A phylogenomic tree of strepsirrhines

Using a phylogenomic dataset comprising 334 nuclear loci with an average length of 3,339 base pairs (bp; range: 158–6,985 bp; total concatenated alignment length: 1,108,850 bp), we reconstructed a phylogenetic tree of Strepsirrhini that includes 71% of all currently recognized species (50% of all lorisiform species and 79% of all lemur species per the taxonomic references in Supplementary Table 1; sample information in Supplementary Data 1). After assessing the impacts of missing data (Supplementary Fig. 1; see Materials and Methods), we used two different species-tree inference approaches (SVDquartets^16^, based on DNA sequence data, and ASTRAL^17^, based on estimated gene trees). Both analyses produced a tree that was largely concordant with prior studies, with Madagascar’s lemurs (infraorders Chiromyiformes and Lemuriformes) as a monophyletic group sister to all strepsirrhine species from Asia and continental Africa (infraorder Lorisiformes) and well-supported clades representing each strepsirrhine family (Fig.1, Supplementary Figs. 2, 3).

**Figure 1.**
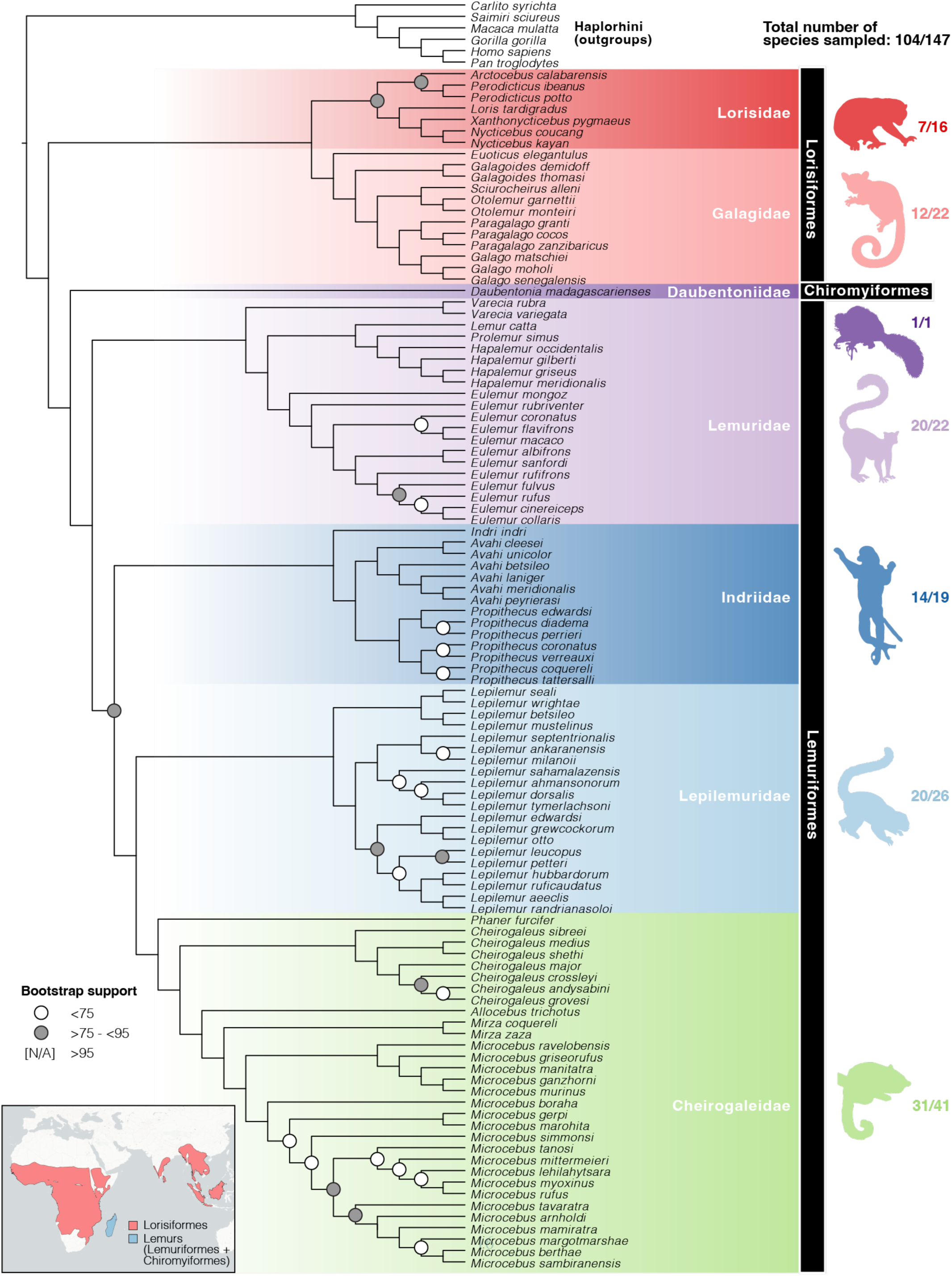
A species tree of strepsirrhine primates estimated using SVDQuartets. Node support values were estimated using 1000 bootstrap replicates. Nodes with > 95% bootstrap support are not labeled, while nodes with 75-95% support and < 75% support are indicated by gray and white circles, respectively. The actual bootstrap support values for all nodes are provided in Supplementary Fig. 2. Branch lengths are not scaled. Infraordinal names are shown in black bars, and family names are shown in different colors matching the silhouette image of a representative species to the right of the tree. Silhouettes were obtained from PhyloPic.org and are credited to T.M. Keesey, R. Lewis, Maky, R.D. Sibaja, and G. Skollar. The numbers next to each silhouette indicate the number of species sampled from each family as a fraction of the total number of described species in the family. Inset map shows the combined distributions of all lorisiform and lemur species in red and blue, respectively. Distribution maps were obtained from the IUCN Red List spatial database.

This nuclear dataset confirms that the family Lorisidae [the ‘slow-climbing’ angwantibos (*Arctocebus*), pottos (*Perodicticus*), and lorises (*Loris*, *Nycticebus*, and *Xanthonycticebus*)] and the family Galagidae [the ‘fast-leaping’ galagos and bushbabies (*Euoticus*, *Galago*, *Galagoides*, *Otolemur*, *Paragalago*, and *Sciurocheirus*)] are reciprocally monophyletic, resolving a longstanding debate (Fig. 1)^10,18–20^. Previous genetic studies have sometimes recovered a sister relationship between galagids and angwantibos/pottos (see also our mitochondrial results below), leading some authors to conclude that traits associated with slow climbing evolved in parallel^10,21^. Although our study does not support parallel evolution, our molecular analyses do recover a relatively short internode (Fig. 2) suggesting that adaptations to slow climbing evolved rapidly.

**Figure 2.**
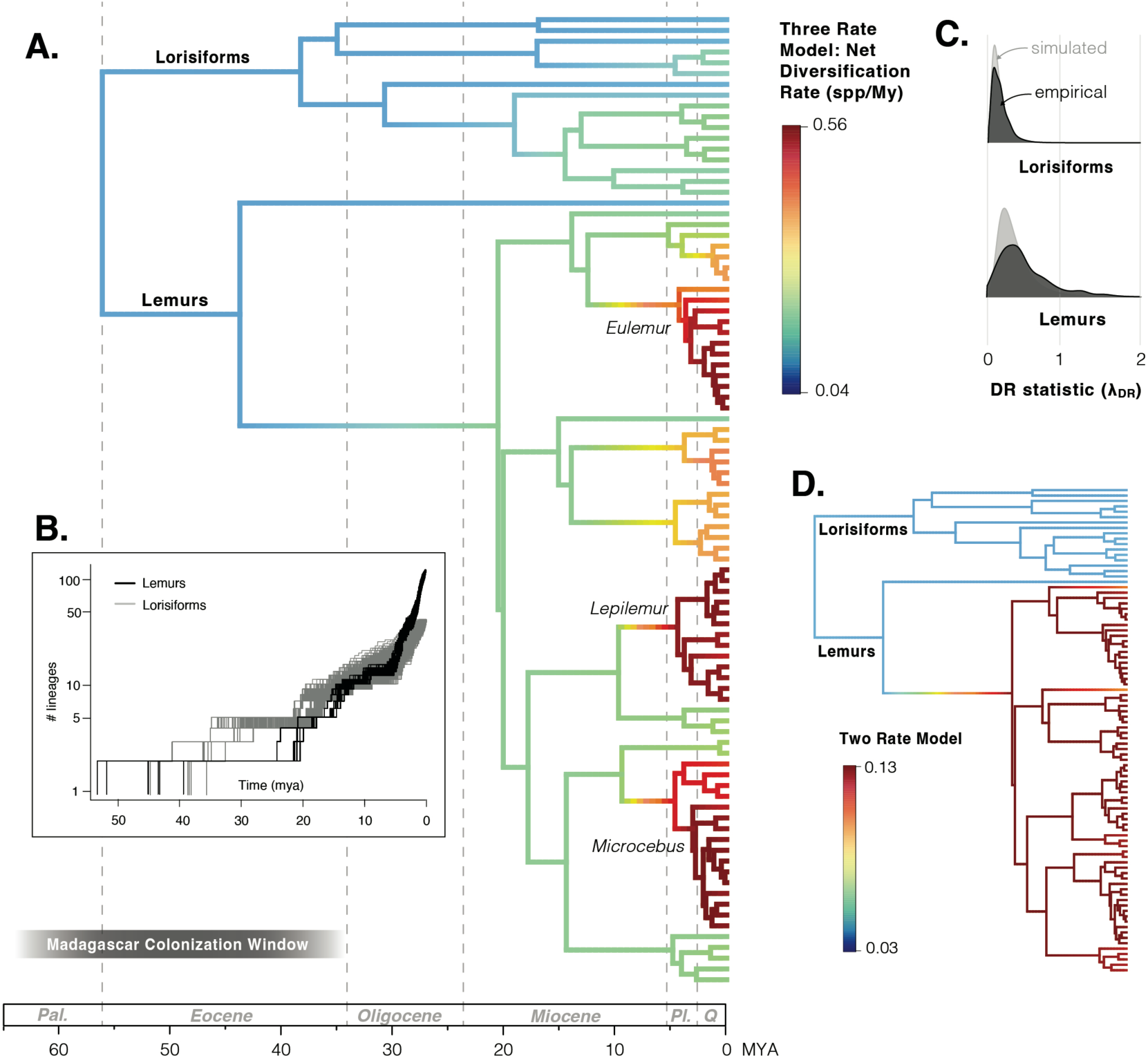
Variation in diversification rate (DR) over time and across strepsirrhine phylogeny. (A) A time-calibrated phylogeny of Strepsirrhini, with branches colored according to DR (species per million years) estimated using a three-rate model in MiSSE. Full results of this analysis can be found in Supplementary Fig. 6. Dashed vertical lines distinguish geological epochs (Pal. = Paleocene, Pl. = Pliocene, Q. = Quaternary). Fully detailed time-calibrated phylogenies showing tip labels, node confidence intervals, and outgroups are shown in Supplementary Fig. 4. Three genera with particularly high speciation rates (*Eulemur*, *Lepilemur*, and *Microcebus*) are indicated on the tree and are discussed in the text. (B) Lineages-through-time plots of lemurs (black) and lorisiforms (gray). Multiple lines are shown to account for six different fossil calibration sets and to account for incomplete taxonomic sampling, which was done by stochastically adding missing taxa to the proper genus to produce a set of 6000 total trees. Note that the y-axis representing the number of lineages is log-transformed. (C) DR variation in lorisiforms and lemurs is visualized as the distribution of tip DR (λ_DR_) values for all species within each taxon. Dark gray distributions represent the empirical λ_DR_ values across the set of 6000 stochastically resolved trees, while light gray distributions represent simulated λ_DR_ values, in which trees of the same species richness as each Family were simulated using a rate-constant birth-death model. (D) MiSSE results, identical to panel A, but estimated using a two-rate model. See Supplementary Fig. 7 for the full visualization of this analysis, and Supplementary Table 5 for MiSSE model testing results.

In lemurs, a major area of phylogenetic disagreement has been the placement of the family Indriidae [the woolly lemurs (*Avahi*), sifakas (*Propithecus*), and indri (*Indri*)]. Some studies have placed indriids as sister to the family Lemuridae [the true lemurs (*Eulemur*), bamboo lemurs (*Hapalemur* and *Prolemur*), ring-tailed lemur (*Lemur*), and ruffed lemurs (*Varecia*)]^22,23^; while other studies have placed indriids as sister to the Cheirogaleidae + Lepilemuridae clade [the mouse lemurs (*Microcebus*), fork-marked lemurs (*Phaner*), dwarf lemurs (*Allocebus*, *Cheirogaleus*, and *Mirza*), and sportive lemurs (*Lepilemur*)]^24–26;^ and still other studies have recovered indriids as the sister group to all of these^27,28^. Our study confirms the placement of Indriidae as sister to Cheirogaleidae + Lepilemuridae, although bootstrap support for this node was 89% (Fig. 1), which was lower than any other family-level relationship on the tree and may be due to an ancient history of introgression (discussed below).

### Timing of the colonization of Madagascar

We dated the strepsirrhine phylogeny using calibrations for nine nodes of the tree based on the fossil record (Fig. 2, Supplementary Table 2)^29,30^. We estimate that lemurs and lorisiforms diverged during the Paleocene or early Eocene [median 55.6 million years ago (MYA), 95% confidence interval (C.I.) = 48.0 – 64.0 MYA] and the crown diversifications of both groups took place during the mid to late Eocene (lemur median = 43.4 MYA, 95% C.I. = 33.9-53.1 MYA; lorisiform median = 38.0 MYA, 95% C.I. = 36.9-40.2 MYA). If we use these 95% C.I.s and assume that lemurs colonized Madagascar from single dispersal event, then the ancestral lemur must have arrived on Madagascar sometime between 33.9 and 64.0 MYA (Fig. 2). However, to account for conflicting recommendations related to fossil placement and node priors among previous studies, we also explored how these times might vary under different fossil calibration regimes (Supplementary Table 2). Across these analyses we observed some conflicting time estimates and large C.I.s on older nodes on the tree (Supplementary Fig. 4), which resulted in a wide window for the timing of the colonization of Madagascar. If we consider the 95% C.I.s across all possible analyses, then lemurs may have colonized Madagascar any time between 26.0 and 82.2 MYA. This window is wider than the full range of possible colonization times estimated by 22 previous studies (Supplementary Table 3), providing a clear example of how sensitive these analyses can be to parameter settings and model choice.

It is worth noting that some recent studies have used mutation rates rather than fossils to calibrate the diversification of mouse lemurs (genus *Microcebus*), and these have produced younger split times than those estimated here. For example, these studies estimated the crown diversification of *Microcebus* at ∼1.5 MYA compared to ∼5 MYA in this study^31–33^. The younger ages recovered in mouse lemur studies may be due to inaccurately elevated pedigree-based mutation rate estimates. Alternatively, our fossil-calibrated tree may overestimate divergence times for young nodes given the dependence on older fossil calibrations deeper in the phylogeny^32^. There are no known lemur fossils on Madagascar that can be used for node calibration, so young lemur nodes are particularly susceptible to overestimation due to reliance on phylogenetically distant fossils. As divergence time estimation is a rapidly changing field, we are hopeful that a consensus may one day be reached using a combination of fossils and demographic modeling. Regardless of the exact timing, it is also important to note that all of these estimates for the colonization of Madagascar assume that all lemurs originated from a single dispersal event. A recent study of African fossils, potentially related to the aye-aye, suggests that Chiromyiformes and Lemuriformes may have colonized Madagascar independently^34^ and we cannot rule out this possibility. If that is the case, then there is much greater uncertainty in the timing of colonization for both groups due to their long stem branches, and further resolution may not be possible at this time.

### Tempo of diversification on Madagascar

Using lineages-through-time plots (Fig. 2b) and Pybus and Harvey’s γ^35^ (a statistic based on internode distances from our ultrametric tree), all of our time-calibrated phylogenies produced clear and significant patterns of increasing diversification rates toward the present in lemurs without a subsequent decline (median γ = 6.57, p-value < 0.0001; Supplementary Fig. 5a). These estimates of γ were still significantly greater than zero even after pruning up to 20 lemur species (median γ = 5.97, p-value < 0.0001; Supplementary Fig. 5a), suggesting that this pattern reflects increasing diversification rates toward the present rather than being an artifact of possible taxonomic inflation^36,37^. To further explore variation in diversification rates across the strepsirrhine tree, we estimated species-specific tip diversification rates (λ_DR_; Supplementary Table 4)^38^. In lorisiforms, we recovered a distribution of λ_DR_ values that tightly mirrored the expectations under a pure-birth model (Fig. 2c). The empirical λ_DR_ distribution for lemurs was also similar to expectations, but included some values that were higher than expected, suggesting that diversification on certain lemur branches cannot be explained by a pure-birth model.

Finally, to visualize variation in macro-evolutionary rates of speciation among branches, we fitted a multi-state speciation and extinction (MiSSE) model which estimates shifts in diversification as a function of one or more hidden states^39^. Across the six time-calibrated trees that we estimated, MiSSE selected models with either two or three hidden states (Supplementary Table 5). All of the two- and three-state models show a clear increase in diversification rate along the branch leading to Lemuriformes (all lemurs except the aye-aye) and all of the three-state models show an additional increase in diversification rate in the last ∼5 million years, which is concentrated on three lemur genera *Microcebus*, *Lepilemur*, and *Eulemur* (clades dominated by red branches on Fig. 2a, Supplementary Figs. 6, 7). The former rate estimates (those along the branch leading to Lemuriformes) should be interpreted with some caution, as this branch is located towards the root of the tree where uncertainty is higher in the state-dependent speciation and extinction models implemented in MiSSE^39^. The estimated rates of diversification towards the tips of the tree (including the elevated rates in *Microcebus*, *Lepilemur*, and *Eulemur*) are comparatively robust.

Lemurs are often cited as a classic example of adaptive radiation, i.e., a clade that diversified from a single common ancestor in response to ecological opportunity. The implied “ecological opportunity” for lemurs was the colonization of Madagascar, an island with presumably underutilized resources at the time of dispersal. The general model for adaptive radiation includes an early burst of rates of speciation and morphological change followed by a decline in both rates as niches become filled – a pattern that has been observed in some Malagasy taxa^40,41^. However, our analyses, as well as some previous studies^6,42^, show that diversification rates in lemurs have not yet declined and may in fact still be increasing. One possible alternative characterization of lemurs is a “constructive radiation”, defined as a radiation that continues to expand as new opportunities are generated continually over time, either due to changing environmental conditions or due to ecological feedback constructed by the radiation itself^43^. ^44^A prediction of constructive radiations is that there may be a lag time between phenotypic disparification and taxonomic diversification^45^. Although we did not assess phenotypic rates of evolution in this study, previous work has shown that morphological disparity in lemurs did evolve quickly after colonization^6^. If we assume that all lemurs (Chiromyiformes + Lemuriformes) originated from a single colonization event 33.9 - 64.0 MYA, then there was a lag time of approximately 10-20 million years before taxonomic diversification rates significantly increased.

It is also important to recognize that many early writers on adaptive radiation did not include explosive speciation as a defining feature at all^46–48^ and it is widely acknowledged that radiations (in the more general use of the term) may arise slowly due to a variety of biotic and abiotic factors, with different predictions for island versus continental radiations^49–51^. Island radiations often begin after an ancestor colonizes a depauperate area, with character displacement among daughter lineages being driven by competition in sympatry. However, on larger continental scales, it is more common for radiations to arise allopatrically as taxa cross geographical barriers or become isolated in habitat fragments when climatic conditions change^52^. Lemurs colonized Madagascar shortly after the Cretaceous-Paleogene extinction event, so they likely arrived on an island that was depauperate and relatively homogeneous in terms of environmental conditions^53^. However, Madagascar is large enough, old enough, and contains enough modern topological and environmental heterogeneity, that speciation has often occurred in allopatry; for example, there are many examples of lemur species boundaries shaped by river barriers, mountains, or watersheds^54–58^. Thus, while Madagascar is an insular, island system, studying Madagascar’s biodiversity only through the lens of island biogeography may overlook patterns that arose through continental processes.

### Recent radiations in Madagascar’s lemurs

In several of the diversification analyses described above, we observed a second burst of diversification in lemurs around the start of the Pliocene (∼5 MYA; Fig. 2a and Supplementary Fig. 6). This pattern is particularly evident in three genera: *Microcebus*, *Lepilemur*, and *Eulemur*. Thus, while lemurs overall might not conform to a traditional definition of adaptive radiation, these three subclades within lemurs might still offer opportunities for scientists who are interested in young, explosive radiations to directly observe ecological speciation, sexual selection, and the spread of key innovations. We suggest that the high diversification rates observed in *Microcebus*, *Lepilemur*, and *Eulemur* around the Miocene-Pliocene transition (or during the Pleistocene, if mutation-date-based divergence times from other studies are applicable^31–33)^ are the result of multiple biotic and abiotic processes. In terms of biotic factors, our results (below) suggest that all three of these genera were experiencing high levels of interspecific gene flow, which might have resulted in novel combinations of alleles that were fuel for diversification. At the same time, the Miocene-Pliocene transition is associated with massive expansions of grasslands and savannas around the world (including Madagascar) as temperatures became cooler^59,60^. The forest ecosystems that had already been established on Madagascar as early as the Cretaceous would have become fragmented during this time, forcing lemur populations into allopatry and contributing to diversification. One additional abiotic factor that may have promoted high speciation rates toward the end of the Miocene is the increasing amount of topological complexity due to montane uplift^61^. The mountains of Madagascar reached their current elevations ∼10 MYA^52^, which could have set the stage for diversification as lemurs adapted to new elevational niches and once-contiguous populations were separated from each other. A line of evidence supporting this hypothesis is that lemur species are tightly linked to specific watersheds that were shaped by these mountains^56^.

### Tempo of diversification in lemurs’ sister group

Previous molecular evidence suggests that lorisiforms have experienced lower rates of diversification relative to lemurs^6^. Our macroevolutionary rate analyses (Fig. 2a, Supplementary Figs. 6, 7) estimated a moderate rise in diversification rate within the family Galagidae (specifically the genera *Galago* and *Paragalago*), but these were still lower overall than most lemuriform clades. These results were concordant with our analyses of λ_DR_, which estimated median values to be more than twice as high in lemurs relative to Lorisiformes: 0.41 species per million years (My) compared to 0.14 species/My, respectively (Fig. 2c, Supplementary Table 4). However, most studies to date have suffered from poor species-level sampling of lorisiforms, and several new species have been identified in recent years, making lorisiform taxonomy and diversification a subject of continuing discussion^10,62,63^. To explore how greater taxonomic attention could influence our estimated rates of diversification, we artificially added up to 20 tips on the lorisiform tree. Interestingly, even with 20 added species (an implausibly high increase in described diversity) lorisiforms would still not match the high rates of diversification seen in lemurs (γ = 5.66 compared to the above-estimated lemur γ = 6.57; Supplementary Fig. 5). That lemurs and lorisiforms have dramatically different rates of diversification is perhaps unsurprising given that lemurs occur singularly on an island, whereas the evolution of lorisiforms has played out over continental scales where several of the niches occupied by lemurs have been occupied by competing species not found on Madagascar (e.g., there are no diurnal lorisiforms, perhaps due to competition with catarrhine primates).

We focused our analyses on extant species because there are no known primate fossils from Madagascar older than the Holocene. An important consequence of using time trees with only extant taxa is that—because lineages that originated recently have had less time to go extinct— there can be a bias toward increased rates of speciation closer to the present^64^. Despite concerns that absolute rates of diversification may be inaccurate in such analyses, previous research suggests that relative differences in state-dependent rates (how fast lemurs radiated relative to lorisiforms, and how quickly specific clades radiated relative to background rates) can still be estimated with high confidence^65^. It is still concerning, however, that time-varying diversification models suffer from non-identifiability; that is to say, an infinite number of speciation and extinction rate functions could be produced from the same phylogeny with equal likelihoods (the so-called congruence class)^64^. To address this specific concern, we used the R package CRABS^66^ to test whether trends in diversification rates remained consistent across models in the congruence class. We evaluated scenarios where extinction rates were (1) initially high but decreased over time, (2) initially low but increased over time, and (3) allowed to fluctuate randomly over time. Remarkably, all models in the congruence class consistently recovered a clear signal of a sudden increase in speciation rate around 5-6 MYA (Supplementary Fig. 8). This suggests that the signature that we recovered of a burst of speciation around the Miocene-Pliocene boundary in lemurs is robust and can be detected even from extant-only time trees. ^65^

### Signatures of introgression in strepsirrhines

Considering the historical difficulty in resolving the strepsirrhine phylogeny, it seems likely that a variety of biological processes, including incomplete lineage sorting (ILS) and introgression, have left signatures in the genome that deviate from the true history of speciation^67^. As a first step toward assessing whether introgression has been present in the evolutionary history of strepsirrhines, we compared the phylogeny generated from our nuclear dataset to a mitochondrial phylogeny (Fig. 3; Supplementary Fig. 9). While we acknowledge that our nuclear dataset is more likely to reflect the true species tree compared to mitochondrial data, topological differences between these two trees can help identify candidate branches of the tree that have experienced introgression^68^. In our case we observed three topological differences between these trees (Fig. 3): (1) Indriidae was sister to all other lemuriform families in the mitochondrial phylogeny, as opposed to Cheirogaleidae + Lepilemuridae in the nuclear phylogeny; (2) the genus *Hapalemur* was sister to *Lemur* in the mitochondrial phylogeny as opposed to *Prolemur*; and (3) in the mitochondrial phylogeny the genera *Perodicticus* and *Arctocebus* formed a clade with galagids, rendering Lorisidae non-monophyletic.

**Figure 3.**
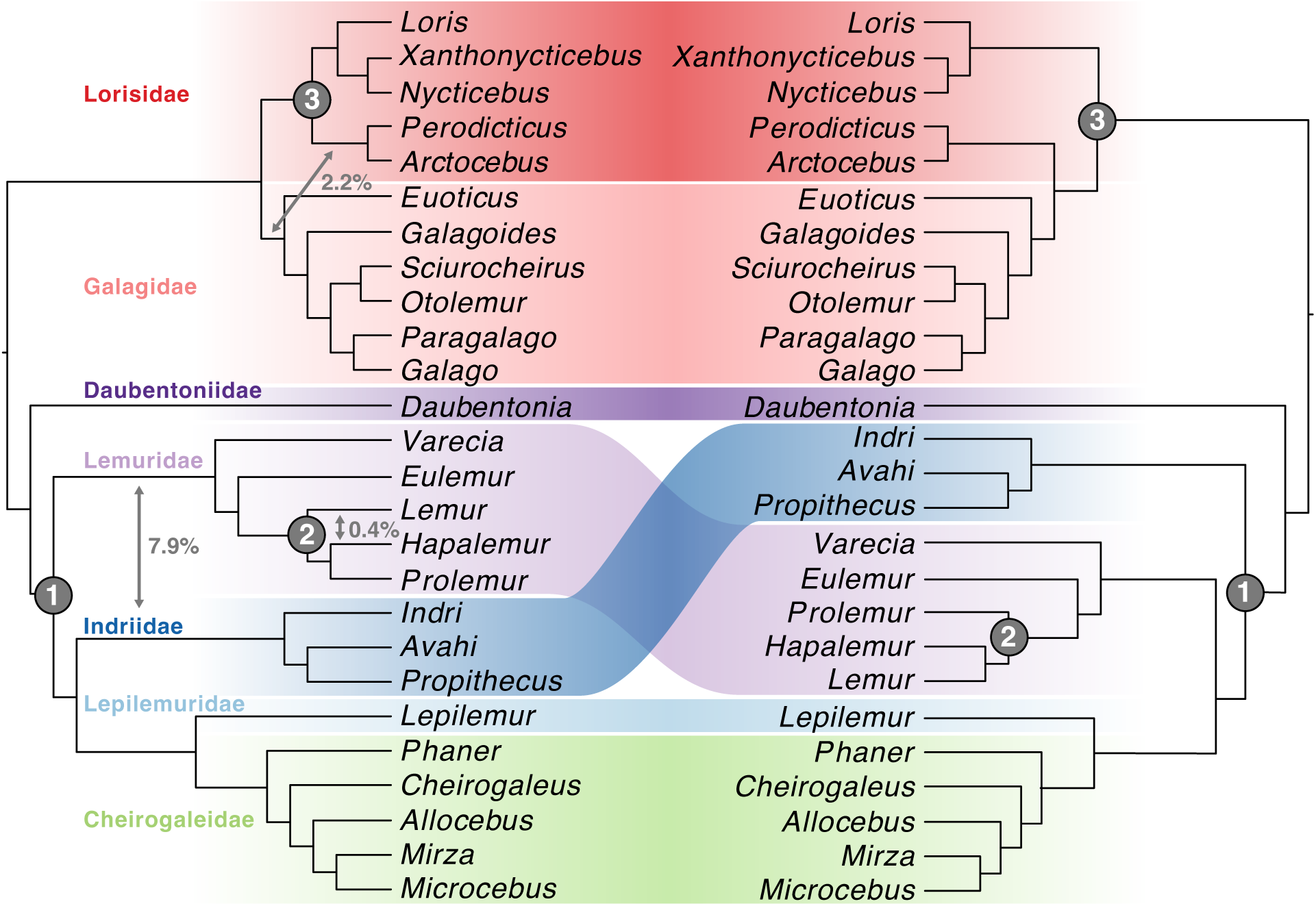
Discordance between nuclear (left) and mitochondrial (right) phylogenies of strepsirrhine genera. Both phylogenies were estimated using IQTree with all available individuals, and then species-level branches were manually collapsed to visualize one branch per genus. Fully detailed phylogenies from these analyses are available in Supplementary Figs. 9 and 13. The nuclear and mitochondrial trees are discordant in three locations labeled with gray numbered circles. Gray arrows on the nuclear phylogeny indicate three locations, where gene flow was inferred using QuIBL, with the proportions of introgressed loci labeled. Families are colored to match Fig. 1.

To explicitly test whether introgression rather than ILS caused these topological differences (labeled with gray circles in Fig. 3) we used the program QuIBL, which estimates the proportion of introgressed loci for each species triplet using gene-tree branch lengths^69^. This analysis recovered significant signatures of introgression for all three topological differences (gray arrows on Fig. 3; full results in Supplementary Fig. 10). On average, we estimated 7.9% introgressed loci for topological difference #1 (between Lemuridae and Indriidae), 0.4% for topological difference #2 (between the genera *Lemur* and *Hapalemur*), and 2.2% for topological difference #3 (between the family Galagidae and the *Perodicticus-Arctocebus* clade). This indicates that an ancient history of introgression likely contributed to topological uncertainty in these regions of the phylogeny. This analysis also identified small but significant proportions of introgressed loci in three other regions of the tree: 0.4% between *Eulemur* and two other lemur genera (*Lemur* and *Varecia*), 0.6% between *Nycticebus* and two other lorisiform genera (*Galago* and *Euoticus*), and 0.7% between lemurs and three lorisiform genera (*Euoticus, Nycticebus,* and *Xanthonycticebus*; Supplementary Fig. 10). These results suggest that introgression was prevalent during early strepsirrhine evolution, likely occurring before and after the colonization of Madagascar and/or among now-extinct lemur relatives in continental Africa.

At shallower taxonomic levels we also observed many uncertain relationships (low or moderate node support) within five lemur genera: *Eulemur*, *Propithecus*, *Lepilemur*, *Cheirogaleus*, and *Microcebus*. One possible reason for low node support could be introgression. To test this hypothesis, we estimated phylogenetic networks for each of the five genera. In all five analyses a model with at least one reticulate branch (*H* = 2–5) was highly supported (Fig. 4, Supplementary Fig. 11). These reticulations are predominantly among ancestral species. However, in two instances we recovered introgression between extant taxa (*Lepilemur tymerlachsoni/L. dorsalis* and *Microcebus lehilahytsara/M. mittermeieri*) which are both parapatric species pairs, making hybridization highly plausible. This suggests that introgression has been prevalent in the evolutionary history of all five of these genera and has likely been an additional source of genealogical conflict beyond ILS. Another recent study also suggested that a burst of speciation (and resulting phylogenetic uncertainty) in the Microcebus clade is the result of an ancient introgressive hybridization event^70^. These results highlight an important consideration for phylogeneticists: if we continue to use species-tree models that only account for ILS as a source of gene tree heterogeneity across the genome, larger genomic datasets will never result in 100% node support for branches affected by a history of introgression.

**Figure 4.**
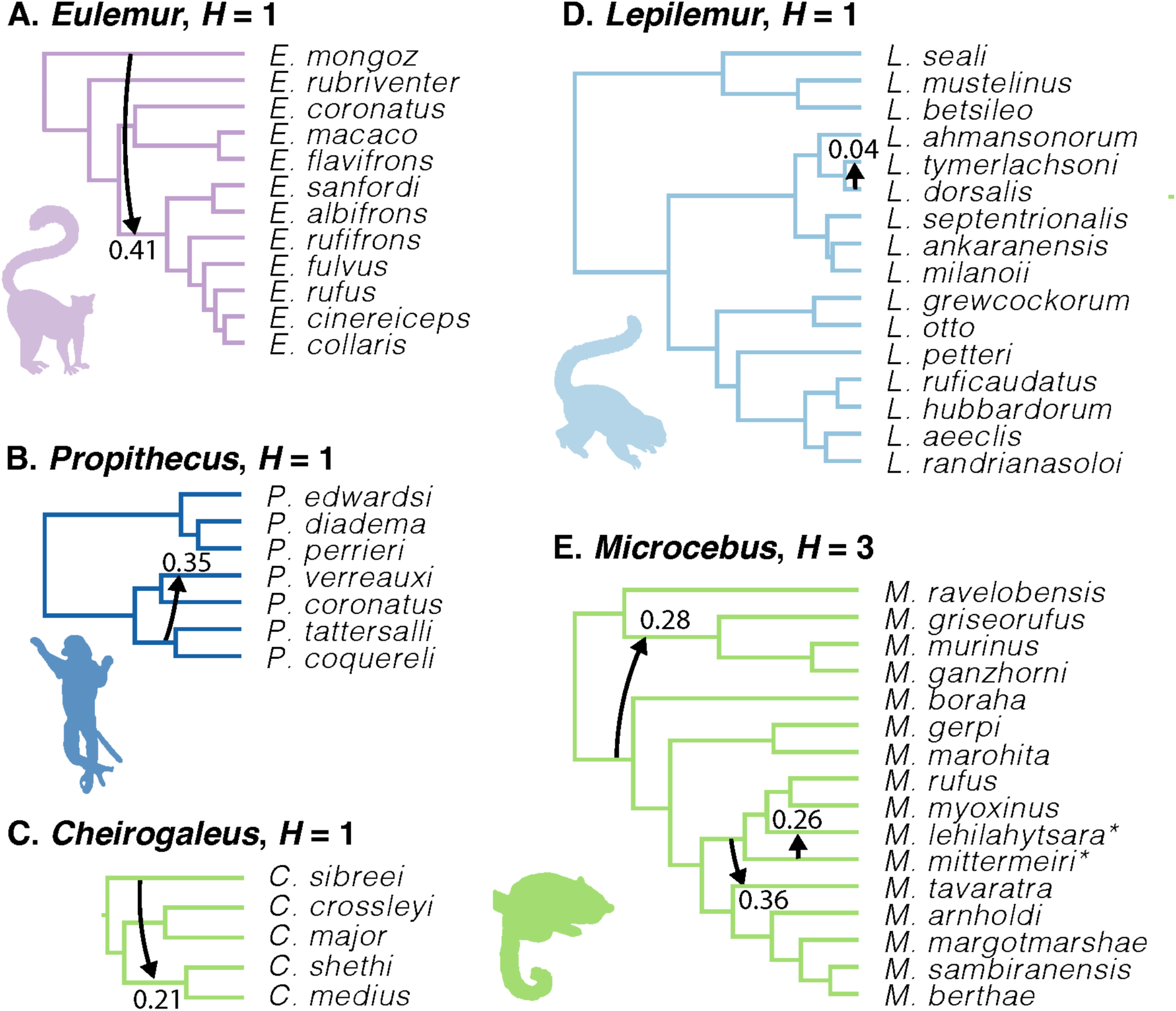
Phylogenies with reticulate relationships estimated by PhyloNet for five strepsirrhine genera (A-E) that had poorly resolved nodes (bootstrap support values < 75%) in our species tree analyses (Fig. 1). Arrows indicate the reticulation events (*H*) and are labeled with the estimated inheritance probabilities. Note that several models received similar support (Supplementary Fig. 11), and here we show the models with the lowest *H* among the well-supported models. Colors and silhouette images for each genus match the Family-level formatting from Fig. 1. *Note that a recent paper ^31^ proposed synonymizing *M. mittermeieri* and *M. lehilahytsara* as a single species; however, we treated these as distinct species as our data did not support a sister relationship.

Gene flow is a topic of keen interest for lemur biologists, as there are several documented active hybrid zones across Madagascar^71^ and introgression appears to have been a staple of lemur evolution on recent timescales^26,72^. Our phylogenomic evidence expands our understanding of the history of hybridization in lemurs by showing that introgressive hybridization is not merely a recent phenomenon but has been a pervasive force throughout the evolutionary history of lemuriforms. Indeed, we identified introgression during the early divergence of families ∼40 MYA (between Lemuridae and Indriidae), during the divergence of genera ∼10 MYA (between *Lemur* and *Hapalemur*), and among species in the same genus within the last ∼5 million years (Figs. 3, 4).

### Hybridizing species experience higher diversification rates

One interesting finding from studies outside of Strepsirrhini has been that some of the most species-rich clades have experienced the highest amounts of introgression^e.g., 15^. To understand whether there is a correlation between diversification rates and introgression in our system, we first scored each species as “hybridizing” or “non-hybridizing”, based on this study as well as an extensive literature review (Supplementary Table 6). We then tested the fit of five different models of diversification (Supplementary Table 7) which varied based on character-dependence or -independence, and on the presence or absence of unsampled factors (“hidden states”); the least complex model (the dull null) assumes a single rate of diversification regardless of whether the taxon hybridizes, whereas the most complex model (the hidden-state speciation and extinction, or HiSSE, model^73^) assumes that diversification rates are influenced both by the presence or absence of hybridization and by hidden states which might be interpreted as noise in the diversification process. We found that the top-ranking model of diversification was a binary-state speciation and extinction (BiSSE) model; i.e., a model in which the diversification rate is tightly correlated with the presence or absence of hybridization without additional hidden states (Supplementary Table 7). Under this state-dependent diversification model, hybridizing species were estimated to have a net diversification rate that was more than four times higher than non-hybridizing species (Table 1). The BiSSE model was supported across multiple variations of this analysis: when lemurs and lorisiforms were analyzed together, when lemurs were analyzed alone, when we treated all unsampled taxa as hybridizing, and when we treated all unsampled taxa as non-hybridizing (Supplementary Table 7).

**Table 1.** Differences in estimated diversification rates (DR) between hybridizing and non-hybridizing taxa, as estimated by a binary state speciation extinction (BiSSE) model. The BiSSE model was the top-ranking model in a HiSSE analysis (Supplementary Table 7).The six models listed below varied in whether or not they included lorisiforms, and in how unsampled taxa were treated by the model.

| **Model** | **DR (Spp/My):**<br>**Non-Hybridizing**<br>**Taxa** | **DR (Spp/My):**<br>**Hybridizing**<br>**Taxa** |
| --- | --- | --- |
| All Strepsirrhines | 8.11 | 40.93 |
| All Strepsirrhines; All unsampled<br>hybridize | 7.22 | 38.66 |
| All Strepsirrhines; No unsampled<br>hybridize | 7.94 | 33.43 |
| Lemurs Only | 10.65 | 46.97 |
| Lemurs only; All unsampled hybridize | 9.65 | 44.73 |
| Lemurs only; No unsampled hybridize | 10.58 | 38.90 |

One caveat of the above results is that taxa were coded using a liberal definition of hybridization, i.e., they were scored as “hybridizing” if hybridization had been directly observed in the wild or captivity, or if any genetic analysis had detected interspecific admixture or gene flow. When we applied a conservative coding scheme, in which species were only coded as “hybridizing” if there was direct documentation of present-day hybrid offspring in the wild, the best-supported models were character-independent (Supplementary Table 7). However, given that many of our results above point to the presence of introgression on ancient evolutionary timescales, we place less weight on these results as we feel that the inclusion of only modern-day hybrids is unrealistically restrictive. One additional caveat is that it may be difficult to disentangle hybridization from taxonomic attention. For example, groups like *Microcebus* that have been featured in multiple species delimitation studies could have higher rates of diversification because more taxa are being split, and these same groups might also have higher rates of hybridization because they are more recently diverged. To explore this idea, we extended the BiSSE models above to fit a multistate speciation extinction (MuSSE) model with two traits: hybridizing vs. non-hybridizing as well as “high taxonomic attention” vs. “low taxonomic attention” (Supplementary Table 6). In this analysis hybridizing taxa still had higher rates of diversification than non-hybridizing taxa, but the magnitude of this difference was impacted by taxonomic attention; specifically, diversification rates were two to four times higher in hybridizing taxa with high taxonomic attention compared to hybridizing taxa with low taxonomic attention (Supplementary Table 8).

Overall, this work contributes to a growing body of evidence that hybridizing species can experience accelerated rates of diversification^13,15^, and it demonstrates that this effect can be amplified by taxonomic attention. These results are in contrast to many other examples across the Tree of Life where hybridization erodes species diversity by replacing high-fitness offspring with poor-fitness hybrids, or by homogenizing gene pools before the speciation process can complete^11^. However, it is important to note that our data do not allow us to disentangle cause and effect of this correlation: is introgression merely a byproduct of rapid speciation, resulting from an insufficient amount of time for reproductive barriers to evolve? Or does hybridization itself drive rapid speciation? In the latter case, one mechanism by which hybridization can promote speciation is through the process of reinforcement, or the accumulation of reproductive barriers through selection against hybrids^14^. Alternatively, a combinatorial view of speciation posits that hybridization might fuel rapid diversification by shuffling old genetic variants or introducing novel alleles to new populations^13,74^. This is a fruitful area of research, and we can point to the three genera (*Eulemur*, *Microcebus*, and *Lepilemur*) that we identified with high diversification rates as well as high levels of introgression, which serve as convenient jumping-off points for future studies.

### Implications for strepsirrhine conservation and future research

Strepsirrhine primates are in the midst of a biodiversity crisis, with approximately 95% of species being threatened with extinction and 90% experiencing population declines^75,76^. From a conservation perspective our findings provide several important advances. First, they point to species and clades that are most prone to hybridization and gene flow. Another recent genomic study found high levels of gene flow in the same lemur taxa we identified^26^, and this result helps explain why lemurs have higher levels of allelic diversity than most other primates despite severe population declines^77,78^. For some taxa gene flow can be a positive force by introducing new genetic variation and adaptive genes, while in others hybridization can lead to genetic swamping and speciation reversal^79^. Conservation practitioners will need to evaluate instances on a case-by-case basis to determine how best to preserve unique genetic variants while also maintaining population sizes, health, and resilience. Second, this study provides a robust phylogenetic framework that future researchers can use to place new species as they continue to be identified. It is worth noting that strepsirrhine taxonomy is a moving target and some groups have received greater taxonomic attention than others^10,80^, which is one reason we tested for the effects of undescribed species and taxonomic biases in this study.

Our results also reiterate that certain branches on the strepsirrhine tree are evolutionarily significant, i.e., lineages that diverged a long time ago and that perform important ecosystem functions, but now contain few living species. Examples include the monotypic genera *Lemur* and *Indri*, and—as an extreme example—the lone member of Chiromyiformes, the aye-aye (*Daubentonia*). Our results show that these lineages are even older than previously recognized and have experienced slower rates of evolution relative to other lemurs. Finally, our study provides a nuanced perspective on the often-neglected lorisiforms, which are difficult to sample and are therefore underrepresented in strepsirrhine research (including the present study, which included 50% taxonomic sampling of lorisiforms compared to 79% of lemur species). A major effort will be needed to understand lorisiform distributions, taxonomy, population sizes, and diversity in the future.

## Methods

### Sampling

We sequenced DNA from 129 individuals obtained as frozen blood or tissues from a variety of sources including museum collections, the Duke Lemur Center, the German Primate Center, and private collections (Data S1). All samples were obtained as part of previous studies or collected under appropriate permits issued by the local governments (e.g., Madagascar, Kenya, Tanzania; Data S1) and approved by ethics boards for animal welfare (IACUC or equivalent) by the institutions involved in the study. Genomic DNA was extracted from frozen tissues using a Qiagen DNEasy Blood and Tissue kit (Qiagen, Inc.) and double-stranded DNA in each extraction was quantified using a Qubit fluorometer (Invitrogen, Inc.). Where DNA quantities were very low, we used a Repli-G whole-genome amplification kit (Qiagen, Inc.) to increase the amount of DNA prior to library preparation. DNA samples were transported to Florida State University to undergo library preparation, Anchored Hybrid Enrichment (AHE)^81^, and sequencing as described below.

### Probe Design for AHE library prep

Briefly, AHE is a DNA sequencing approach that is widely used in phylogenomics because it targets thousands of conserved protein-coding exons (and their more variable flanking regions) across the taxa of interest. To generate an AHE probe set for strepsirrhines, we adapted the Amniote 2 AHE design^82^ using six previously published genomes: *Daubentonia madagascariensis* (Daubentoniidae, NCBI accession GCA_000241425.1), *Microcebus murinus* (Cheirogaleidae, NCBI accession GCA_000165445.3), *Propithecus coquereli* (Indriidae, NCBI accession GCA_000956105.1), *Eulemur flavifrons* (Lemuridae, GCA_001262665.1), *Eulemur macaco* (Lemuridae, ncbi accession GCA_001262655.1), and *Otolemur garnettii* (Galagidae, GCA_000181295.3). Using the methods/scripts of Hamilton et al.^83^, we identified sequence regions in the strepsirrhine genomes that were homologous to the Vertebrate AHE probe regions developed in Lemmon et al.^81^, extracted 6000bp regions containing those homologs, aligned the sequences across the six strepsirrhine sequences for each locus, then trimmed those alignments to retain only well-aligned regions. After reducing the resulting alignments to a set that had no overlapping regions (some loci in the Amniote kit were from neighboring exons that overlapped when extended), we masked repetitive regions ^see^ ^83^. The resulting alignment covered ∼1.3Mb. We tiled 120bp probes across all remaining sequences at 2.8x density to produce 110809 sequences.

### Library preparation and DNA sequencing

We prepared and sequenced libraries using the AHE protocol, following Lemmon et al.^81^ and Prum et al.^84^. Extracted DNA was sonicated to 250-500bp using a Covaris Ultrasonicator in 96-well glass plates. We performed blunt-end repair and Illumina adapters ligation (with 8bp indexes) using a Beckman Coulter FXp liquid-handling robot. The prepared libraries were pooled in groups of 24 samples, then enriched using an Agilent Sure Design XP kit containing the probes described above. Enriched libraries were pooled, assessed for quality via Bioanalyzer and qPCR (using a Library Quantification Kit from KAPA Biosystems, Inc.), then sequenced and average of 10.9 million read pairs per sample on an Illumina NovaSeq 6000 instrument using paired-end 150-bp chemistry. Sequencing was performed at the Translational Lab in the College of Medicine at Florida State University.

### Retrieval of DNA sequence data from previously published genomes

We supplemented our sampling with previously published whole genome data from six primate outgroups (from UCSC genome browser: human-hg38, chimpanzee-panTro6, gorilla-gorGor5, rhesus-rheMac8, squirrel monkey-saiBol1, tarsier-tarSyr2) and 18 previously published strepsirrhines (Supplementary Table 9). We mapped probe region sequences from the lorisiforms probe design alignments (see above) and extracted the matching sequences from each downloaded genome.

### Quality control and AHE Assembly

Newly sequenced reads were demultiplexed and quality filtered using Casava (Illumina, Inc.). Quality-filtered Illumina reads were merged following a custom bioinformatic pipeline outlined in Rokyta et al.^85^. This process resulted in merged reads that had sequencing adapters removed, and sequencing errors corrected. Following Hamilton et al.^83^, we assembled the reads using a quasi-*de novo* approach where the strepsirrhine probe region sequences were used as references for assembly. The resulting consensus sequences were filtered, with those that resulted from at least 83x read depth being kept for downstream analyses. We performed orthology across the consensus sequences (and genome-derived sequences mentioned above) using a neighbor-joining approach to identify a single othologous sequence per individual at each AHE locus (see Hamilton et al.^83^ for details).

### DNA alignment

Sequences determined to be orthologous were aligned using MAFFT (v7.023b) ^86^, then trimmed/masked using default settings (see Hamilton et al.^83^ for details). As a last step, all loci were imported into the software Geneious v.2022.2 ^87^ for a final quality check by eye, with poorly aligned regions being fixed using the Local Realignment tool. Final alignments for each locus were exported from Geneious to nexus, phylip, and fasta files for further analysis.

### Evaluation of the effects of missing data

To evaluate the impact of missing data on phylogenetic analyses, we first created six concatenated fasta files containing:

(1) All loci, all individuals (161 individuals, 1,108,850 bp)
(2) All loci, individuals with > 50% missing data removed (144 individuals, 1,108,850 bp)
(3) All loci, individuals with > 20% missing data removed (106 individuals, 1,108,850 bp; outgroups with > 20% missing data were retained for rooting)
(4) Reduced loci (dropping 37 loci that failed to sequence in lorisiforms and outgroups), all individuals (161 individuals, 969,767 bp)
(5) Reduced loci, individuals with > 50% missing data removed (144 individuals, 969,767 bp)
(6) Reduced loci, individuals with > 20% missing data removed (109 individuals, 969,767 bp; outgroups with > 20% missing data were retained for rooting)

All six datasets were analyzed using IQ-TREE v.2.1.3 ^88^. Each locus was treated as a separate partition for automatically estimating substitution models, and a maximum-likelihood phylogeny was estimated for each dataset with 1000 ultrafast bootstrap replicates.

We observed that missing data had no effect on the overall topology or node bootstrap support values, except that several important genera and species were removed from the datasets with reduced taxa. However, we observed that many of the taxa with > 50% missing data had long terminal branch lengths (Supplementary Figs. 12-17). Because branch lengths are important in diversification and divergence time analyses, we ran our divergence time analysis (see below) using Dataset 2 (all loci, taxa with > 50% missing data removed).

### Phylogenetic analysis

We estimated species trees using two different approaches: (1) an alignment-based analysis in the program SVDquartets ^16^ and (2) a gene-tree-based analysis in the program ASTRAL ^17^ Both are coalescent programs that use quartet scores to select the best species-tree topology. We ran SVDquartets in PAUP* ^89^ using a concatenated sequence file as input. We used multilocus bootstrapping and the evalq=all setting, which specifies that all quartets should be evaluated, and designated the five haplorrhine species as the outgroup. Finally, we ran ASTRAL with default settings using individual gene trees from each locus as input. These gene trees were generated from individual sequence alignments for each locus using RAxML-ng ^90^ under the GTR model.

### Estimation of divergence times

We estimated time-calibrated phylogenies using the MCMCTree algorithm^91^, implemented within the program PAML^92^. We used our SVDquartets topology and full concatenated dataset as the inputs for this analysis, but pruned the input files to include only taxa with < 50% missing data and only one individual per species (the individual with the lowest proportion of missing data). Divergence time estimation was performed six times using different fossil calibration sets based on recommendations from previous studies (Supplementary Table 2). Prior distributions on these nodes were visualized using the R package MCMCtreeR^93^. We also used the R package ddBD^94^ to estimate the parameters for the birth-death model from our empirical data using the sum of squared errors method for selecting the initial values in grid search (BDparas = 21.011 17.752 0.71). We used the GTR+G model (model = 7) with 5 gamma categories (ncatG = 5), and used the program baseml (distributed with PAML) to estimate the alpha parameter (alpha = 0.40337) and substitution rate (rgene_gamma = 1 14.2282). We ran the MCMCTree analysis using an approximated likelihood approach,^95^ where the gradient and Hessian of the likelihood function are estimated first (usedata = 3), then divergence times are estimated using Markov Chain Monte Carlo (MCMC; usedata = 2). The first 20,000 iterations of the MCMC were discarded as burn-in, then we ran the MCMC chain for 1 million iterations sampling every 20 for a total of 50,000 samples.

### Macroevolutionary rates of speciation

To generate lineages-through-time (LTT) plots, we used our time-calibrated phylogenies as input to the ltt function in the R package phytools ^96^ To visualize potential variation in these plots that might be caused by incomplete taxonomic sampling, we also generated a suite of 1000 trees for each of our six time-calibrated phylogenies using the program TACT ^97^, to stochastically add all missing species to the proper genera, and estimated an LTT plot for each of the 6000 stochastically resolved trees. Finally, we used the mccr function in phytools to estimate Pybus and Harvey’s γ [a metric that uses internode distances on an ultrametric tree to infer whether accelerations in diversification rate occurred early (negative γ) or late (positive γ) in the phylogeny]^35^ using the rho parameter to account for sampling fraction.

Some studies have suggested that lemurs are taxonomically over-split.^36,37^ To test whether the results above would be robust to taxonomic synonymization, we wrote a custom R script using commands from the phytools package^96^ to randomly drop five lemur species from each of our 6000 time-calibrated and stochastically resolved phylogenies, leaving at least one representative from every genus. This process was then repeated with 10, 15, and 20 lemur species dropped from the trees. We estimated Pybus and Harvey’s γ for each tree and visualized the distribution of γ values at every level of taxonomic synonymization using ggplot2. It is also possible that lorisiform diversity is *underestimated* due to lack of taxonomic attention; thus, we conducted the same analysis described above, but instead of randomly dropping tips we randomly added five, 10, 15, or 20 “new” lorisiform species on each tree. The new tips were added to regions of the tree < 10 million years old, as we felt that it was unlikely that very ancient lineages have not yet been discovered.

Macroevolutionary rates were estimated using the missing state speciation and extinction model (MiSSE) ^39^, which belongs to the speciation and extinction family of models ^e.g., 73,98–100^ and reconstructs diversification rates as a function of one or more hidden states. We performed this analysis on each of our six time-calibrated phylogenies in R using the package hisse ^73^, setting the estimated proportion of extant species sampled in the phylogeny (*f*) to 0.71^73^. We tested five models which varied in the number of hidden states from one to five, each with an associated turnover rate and extinction fraction. The top-ranking model was selected using the Akaike Information Criterion (Supplementary Table 5) and this model was then used to reconstruct rates across the trees using the function MarginReconMiSSE (Supplementary Figs. 6,7).

We also estimated tip diversification rates (λ_DR_; Supplementary Table 4), which reflect the weighted inverse of phylogenetic branch lengths leading to each tip^38^. A median λ_DR_ value was calculated for each tip across all 6000 phylogenies that were estimated previously using TACT. We also simulated λ_DR_ distributions expected under a homogeneous birth-death process in order to identify specific regions of the tree with higher or lower empirical speciation rates than expected, following the procedure outlined in Upham et al.^101^. Tree simulations and calculation of λ_DR_ metrics were performed using a custom R code (https://github.com/keverson25/StrepsirrhineAHEs/blob/main/TipDR_Calculation.R).

### Comparison of mitochondrial and nuclear datasets

Mitochondrial sequences were captured as off-target reads in our sequencing protocol and were harvested from our raw sequence data (forward and reverse fastq.gz files) using the program MitoZ with the “Chordata” clade setting ^102^ This pipeline retrieved mitochondrial sequence data in 104 individuals. Sequences were assembled and aligned using Geneious ^87^ and a mitochondrial phylogeny was estimated using the IQ-TREE web server ^103^ with default settings, allowing the substitution model to be ascertained automatically (Supplementary Fig. 9). Genera were collapsed into single branches in the main text for visualization purposes (Fig. 3).

### Tests of hybridization and gene flow

We used the program QuIBL^69^ to distinguish between ILS and ancient introgression (above the species level) in our dataset. We first calculated a set of gene trees (one for each locus), using RaxML-ng^90^, with the ultrafast bootstrapping method and the GTR model of substitution. To prepare these gene trees for QuIBL, we used a custom R script to collapse each genus into a single tip and retained only the gene trees with all genera present. We evaluated all triplets and set the outgroup of our species tree to the Haplorhini. We set the ‘numdistributions’ parameter to 2, which corresponds to one branch-length distribution for ILS and one for introgression, and we used default recommendations for the remaining parameters (likelihoodthresh, numsteps, and gradascentscalar).

We also used the program PhyloNet ^104^ to test for introgression in each genus that had topological uncertainty (low node support) in our species-tree analyses (i.e., *Cheirogaleus, Eulemur*, *Lepilemur*, *Microcebus*, and *Propithecus*).^90^ The set of gene trees that was previously estimated in RaxML-ng was used as input to PhyloNet, using a custom R script to prune each gene tree to include only the species from the genus of interest. We used the maximum pseudo-likelihood approach to estimate quartet counts under models that varied in the number of hybridization events (*H*), which we allowed to vary from zero to five. To choose the correct *H* for each analysis, we visualized the log-likelihood scores for each analysis and used the lowest value of *H*, beyond which little improvement in likelihood was observed (Supplementary Fig. 11).

To understand whether there was a correlation between speciation rates and hybridization, we first scored each strepsirrhine species as “hybridizing” or “non-hybridizing” based on this study as well as an extensive literature review using the search engine Google Scholar, where the species name was paired with the words “hybrid” and “introgress” and their structural variants (e.g., “hybridize” and “introgression”; Supplementary Table 6). We generated two scoring systems: (1) a conservative system, where species were only classified as “hybridizing” if there was documentation of that species hybridizing in the wild, and (2) a liberal system, where species were classified as “hybridizing” if there was any documentation of that species hybridizing in the wild or captivity, or if any previous study had found evidence of gene flow using phylogenetic or population genetic analyses, or if that species was a descendant of a reticulate branch leading to one or two tips in our PhyloNet analyses (Fig. 4). Because we were concerned that hybridization might be artificially inflated in the genera where we explicitly looked for evidence of hybridization (i.e., genera that we included in PhyloNet), we also conducted a second round of PhyloNet analyses where we scanned each Family following the same procedure outlined above. These analyses did not reveal any additional taxa to score as “hybridizing” in the liberal system (Supplementary Fig. 18). After finalizing our scoring system, we used an approach similar to Patton et al.^15^, who applied the hidden-state speciation and extinction (HiSSE) trait-dependent diversification model^73^. We evaluated a total of five competing models using the hisse package in R ^73^, ranging from a null character-independent model with a single diversification rate, to a full character-dependent HiSSE model accounting for hidden states. Parameter values were estimated from the top-ranking models, which were selected using the Akaike Information Criterion. We applied this model-testing framework to both the liberal and conservative coding schemes. Then, to understand the influence of unsampled taxa where hybridization status is unknown, we re-ran those tests again using two different values for the sampling parameter *f*: one in which all unsampled taxa on the phylogeny are assumed to hybridize, and one in which all unsampled taxa on the phylogeny are assumed not to hybridize.

Where the top-scoring model was a BiSSE model (see results), we explored how these results were affected by taxonomic attention. To do this we coded taxa as “high taxonomic attention” or “low taxonomic attention” by searching Academic Search Complete (EBSCO Industries, Inc.) for peer-reviewed articles with “[genus name]” in the title and any of the words “evolution”, “phylogeny”, or “population genetics” in the article contents. All members of genera with less than five article hits were coded as “low taxonomic attention” (n genera = 13) while those with five or more hits were coded as “high taxonomic attention” (n genera = 13). This trait was then used in conjunction with the “hybridizing/non-hybridizing” trait to fit a multi-state speciation extinction (MuSSE) model using the R package hisse ^73^.

### Data Availability Statement

All DNA sequence data have been deposited in the NCBI SRA under BioProject ID PRJNA957840. Analytical files have been deposited on FigShare (https://figshare.com/s/790074613c764d6c8cd7).

### Code Availability Statement

Custom R scripts used in this manuscript are available on GitHub (https://github.com/keverson25/StrepsirrhineAHEs).

### Inclusion and Ethics Statement

Samples were obtained from a wide variety of sources (see Supplementary Data S1) including field sampling. Local and international ethical guidelines were followed to minimize disturbance to animals and the environment. Approvals were granted by Madagascar National Parks, the Ministére de l’Environmnement et du Développement Durable de Madagascar and the Committee for Environmental Research (permit numbers 004-MEF/SG/DGEF/DADF/SCB, 072-MINENV.EF/SG/DGEF/DADF/SCB, 100-MINENV.EF/SG/DGEF/DPB/SCBLF, 124/09/MEFT/SG/DGEF/DSAP/SLRSE, 130/16/MEEF/SG/DGF/DAPT/SCBT.Re, 137/13/MEF/SG/DGF/DCB.SAP/SCB, 186/11/MEF/SG/DGF/DCB.SAP/SCB, 78/17/MEEF/SG/DGF/DSAP/SCB.Re, 79/17/MEEF/SG/DGF/DSAP/SCB.Re, 82/18/MEEF/SG/DGF/DSAP/SCB.Re). Samples were exported under CITES permit 19US36412D/9 and imported to the U.S. under U.S. Fish and Wildlife Service permit numbers 2019NW2505894-905. Capture and handling procedures followed routine protocols approved by the Institute of Zoology, University of Veterinary Medicine Hannover Foundation. In the main text, we include a discussion of potential implications of our research on conservation efforts of endangered species.

## Acknowledgments

In memoriam of Elke Zimmermann, who was an esteemed and influential collaborator in the early stages of this project. We also recognize the mentorship, support, and collaboration of the late Judith C. Masters. We thank the following individuals for their lab, field, and infrastructure support: B. Allen, B. Andriatsitohaina, J. Cherry, M. Craul, N. Daniel, I. Dröscher, E. Ehmke, M. Rina Evasoa, M. Foley, K. Freeman, A. Greven, A. Hasainaina, Z. Hert, B. Iambana, A. Jones, A. Junge, A. Katz, C. Kerrick, G. Kett, F. Kiene, M. Kortyna, R. Lewis, M. Le, M. Matocq, R. McGinnis, R. Munds, H. Rafalinirina, T. Rakotonanahary, R. Rakotondravony, M. Ramsay, F. Rasambainarivo, O. Schülke, H. Teixeira, D. Weisenbeck, C. Welch, C. Williams, and S. Zehr. We are grateful to the following museum curators and collection managers for their permissions and assistance with sample acquisition: S. Schaefer and S. Katanova (Ambrose Monell Cryo Collection, American Museum of Natural History); C. Phillips and H. Garner (Natural Science Research Laboratory, Museum of Texas Tech University); N. Duncan and G. Amato (Department of Mammalogy, American Museum of Natural History); and E. Gilissen (Mammals Collection, Royal Museum for Central Africa). Finally, we thank the following organizations for facilitating fieldwork and logistical assistance: the Madagascar Biodiversity Project Field Team, Ankoatsifaka Research Station, Betampona Strict Nature Reserve, Centre ValBio (CVB), Ministère de l’Environnement et Developpement Durable, Madagascar National Parks (MNP), Madagascar Fauna Group (MFG), Madagascar Institute for the Conservation of Tropical Ecosystems (MICET), Groupe d’Etude et de Recherche sur les Primates de Madagascar (GERP), Planet Madagascar, Système des Aires Protégées, and the Committee for Flora and Fauna (CAFF/CORE).

## Funding

U.S. National Science Foundation grant DEB-1355000 (ADY, DWW)

U.S. National Science Foundation grant DEB-2207198 (KME, DWW)

U.S. National Science Foundation grant BCS-1926215 (MEB)

U.S. National Science Foundation grant BCS-1926105 (LP)

U.S. National Science Foundation grant OIA-1826801 (KME)

University Research Postdoctoral Fellowship, University of Kentucky (KME)

Groupe d’Étude et de Recherche sur les Primates de Madagascar (UR)

Otto-Stiftung (UR)

German Federal Ministry of Education and Research (Bundesministerium für Bildung und Forschung) grant 01LC1617A (UR)

German Research Foundation grant DFG Ra502/7-1 (UR)

German Research Foundation grant DFG Ra502/20-1 (UR)

Saint Louis Zoo Field Conservation for Research Fund (MAB)

Duke University Center for International Studies (MAB)

Duke Graduate School (MAB)

Nicholas School of the Environment (MAB)

## Author contributions

Conceptualization: KME, LP, DWW, ADY

Formal analysis: KME, CJP, RZ

Resources: LP, MAB, MEB, MED, PMK, ACK, ARL, EML, UR, BR, RMR, SR, CR, DZ, JS, ADY, DZ, DWW

Writing – Original Draft: KME, LP, DWW

Writing – Review & Editing: MAB, MEB, MED, PMK, ACK, ARL, EML, CJP, UR, BR, RMR, SR, CR, JS, ADY, RZ, DZ

Supervision: LP, DWW

Funding Acquisition: KME, MAB, LP, UR, ADY, DWW

## Additional Information

Supplementary Information is available for this paper.

Correspondence and requests for materials should be addressed to Kathryn M. Everson.

