## Supplementary figures and images for "Multiple bursts of speciation in Madagascar’s endangered lemurs"

### Supplementary Fig. 1

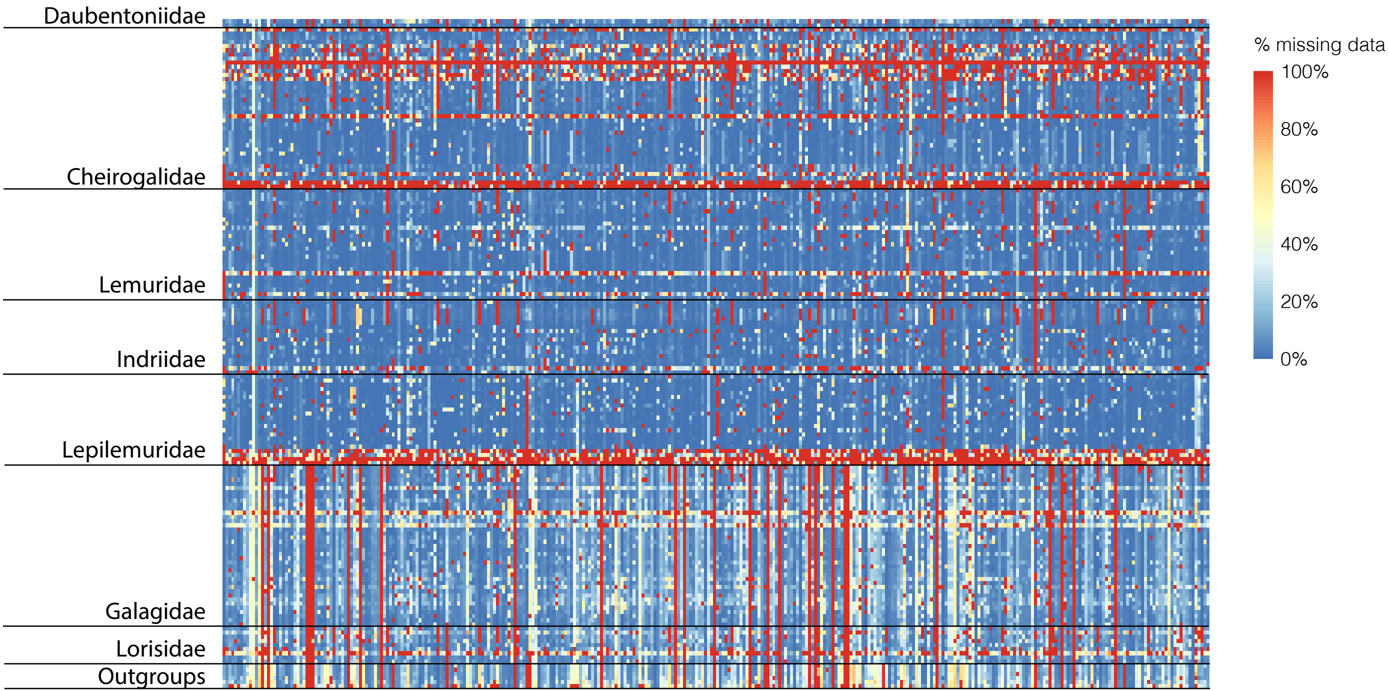

### Supplementary Fig. 2

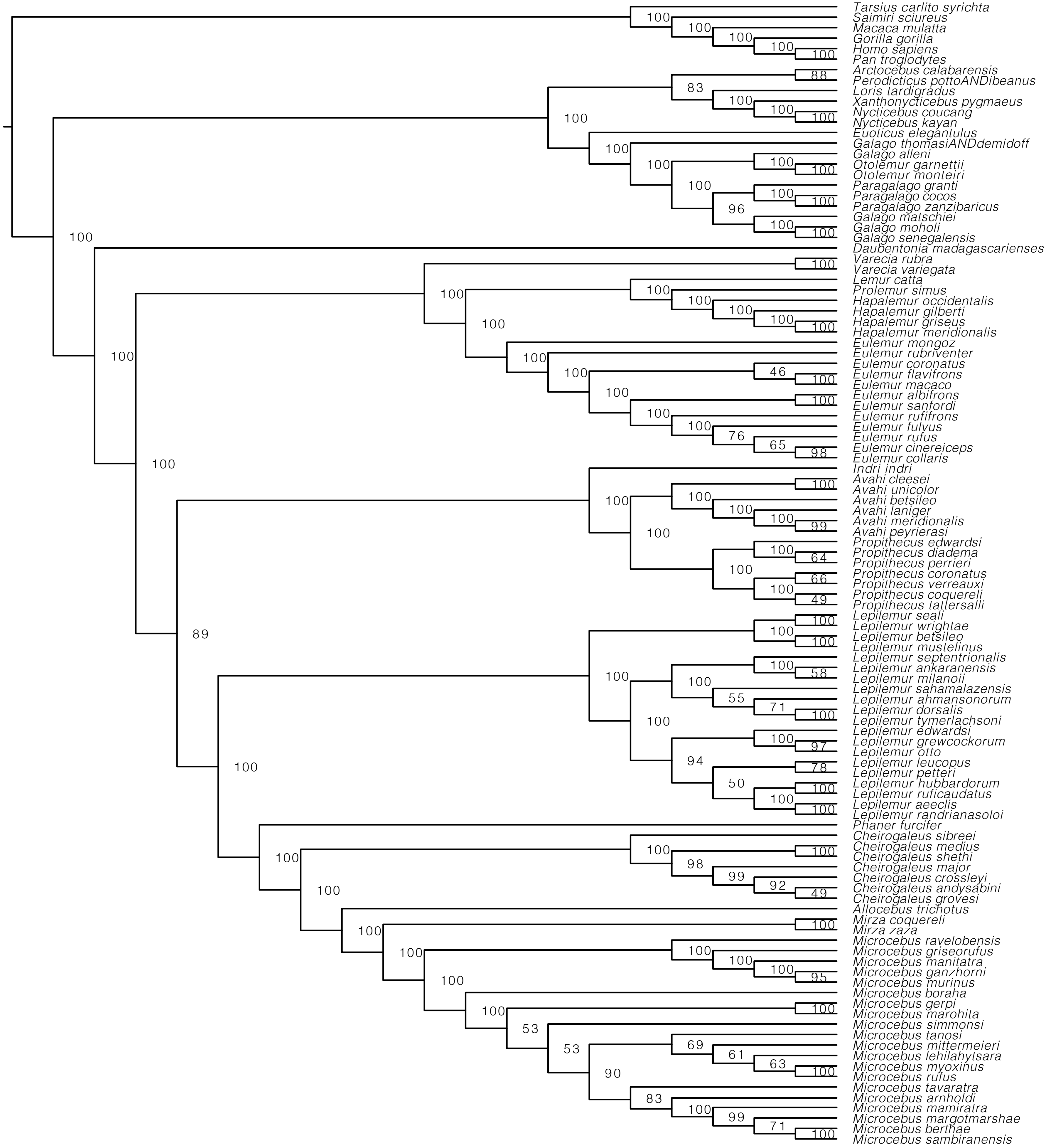

### Supplementary Fig. 3

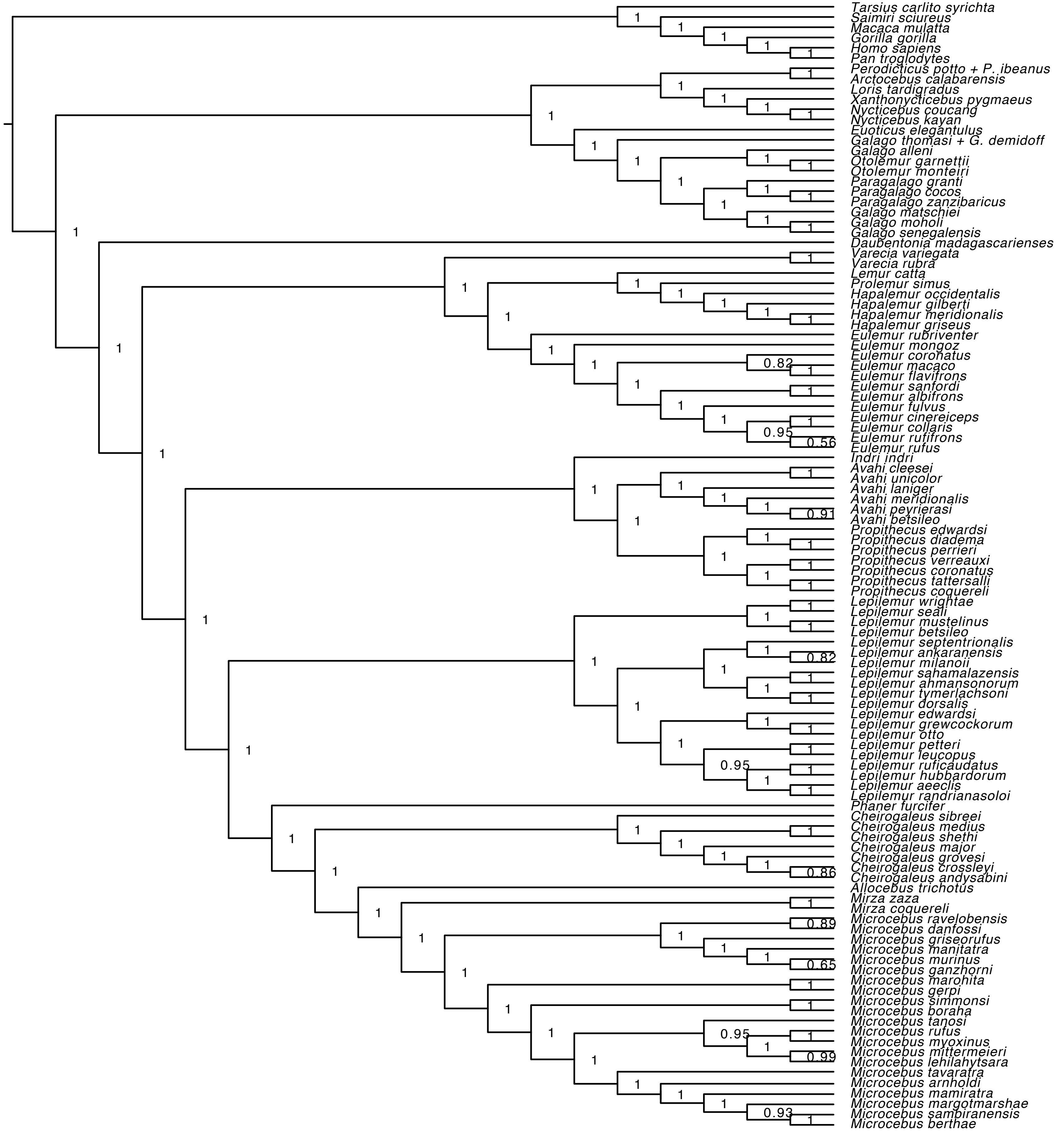

### Supplementary Fig. 4

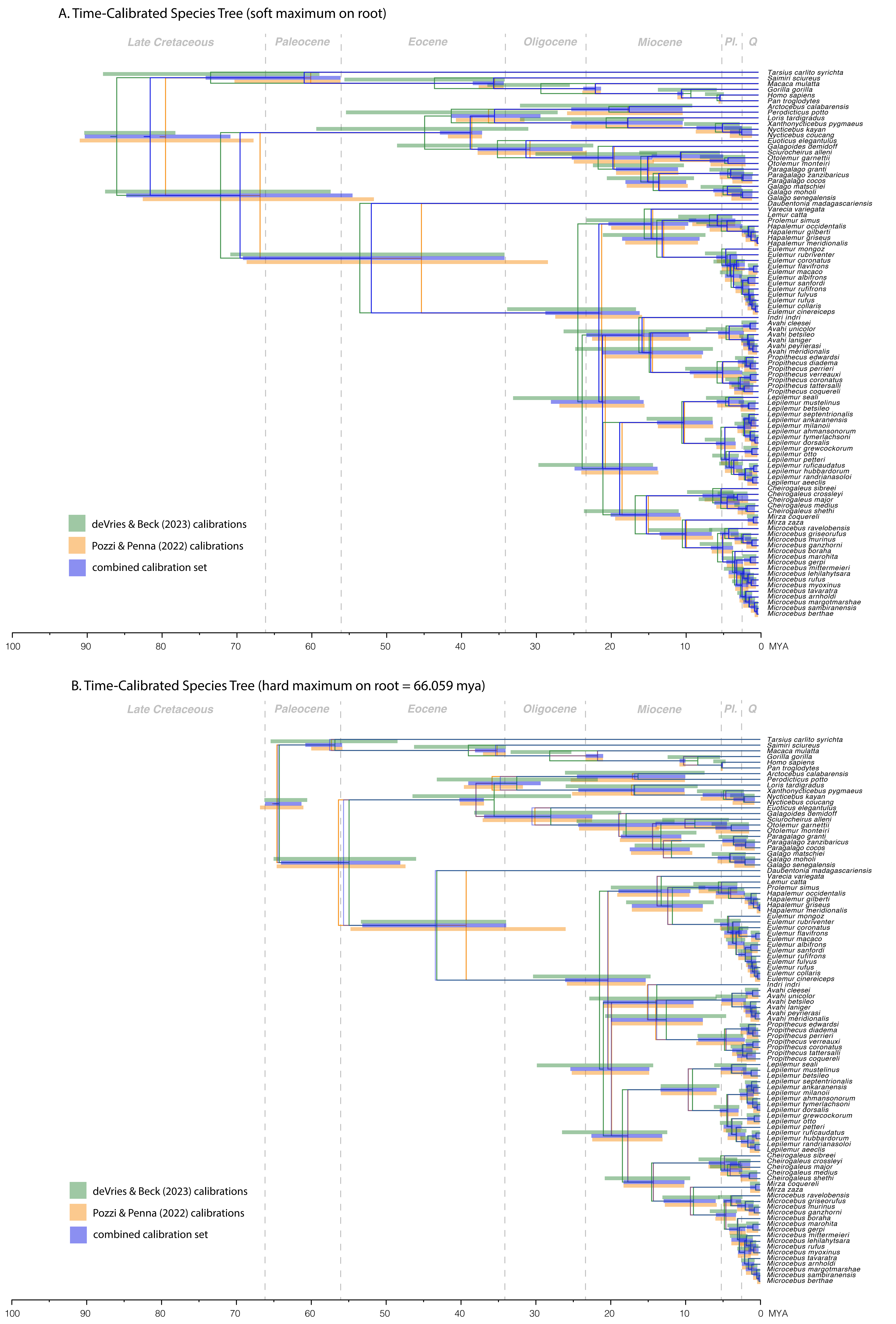

### Supplementary Fig. 5

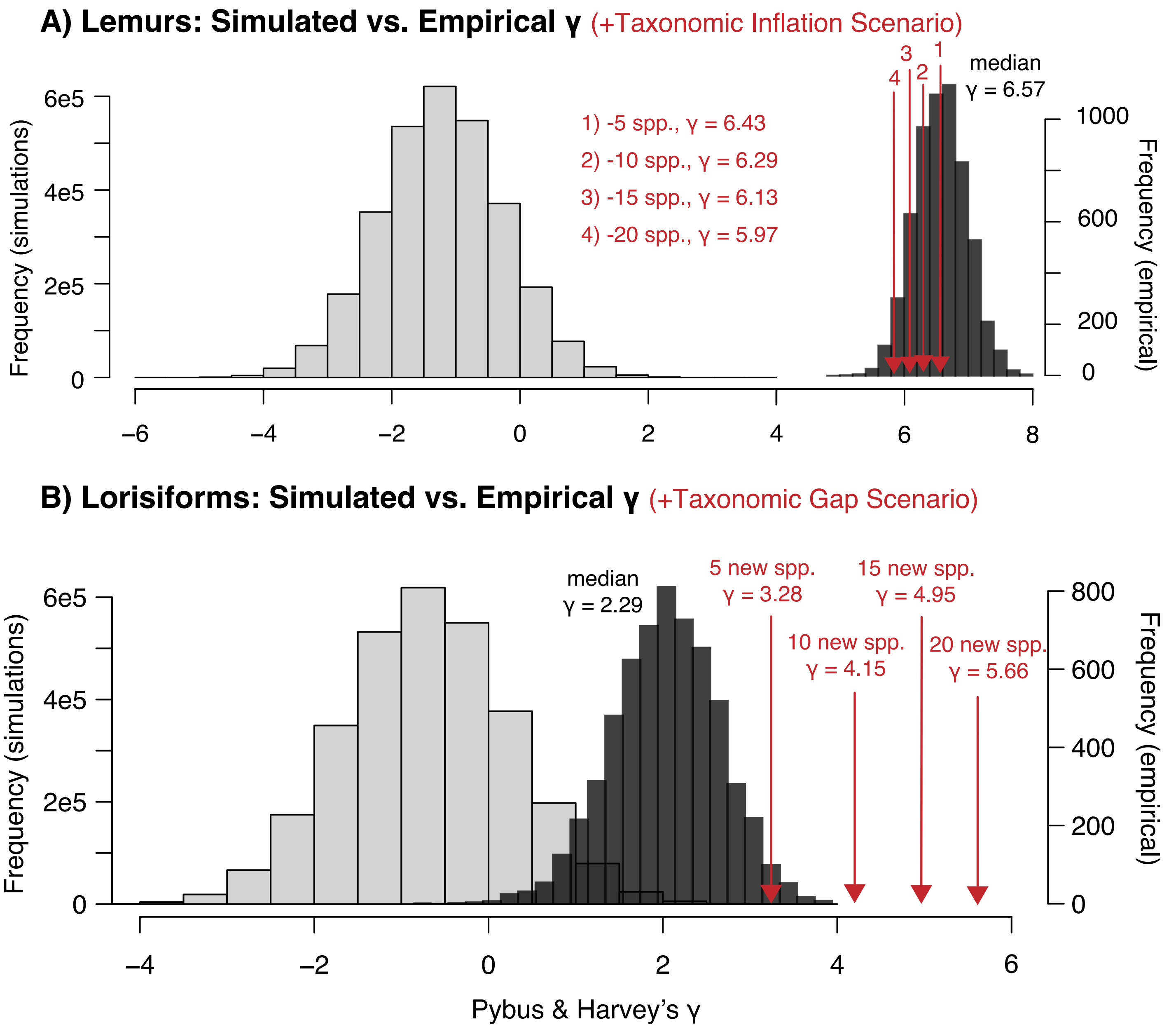

### Supplementary Fig. 7

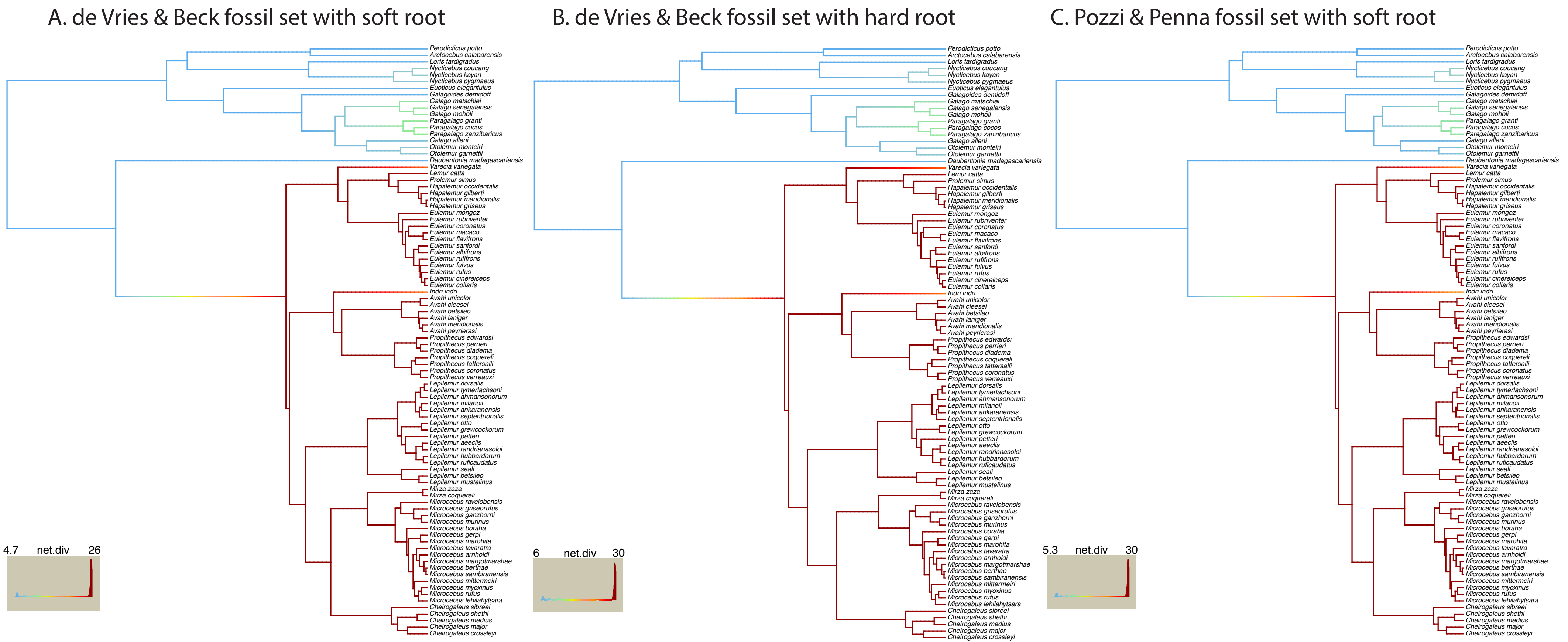

### Supplementary Fig. 8

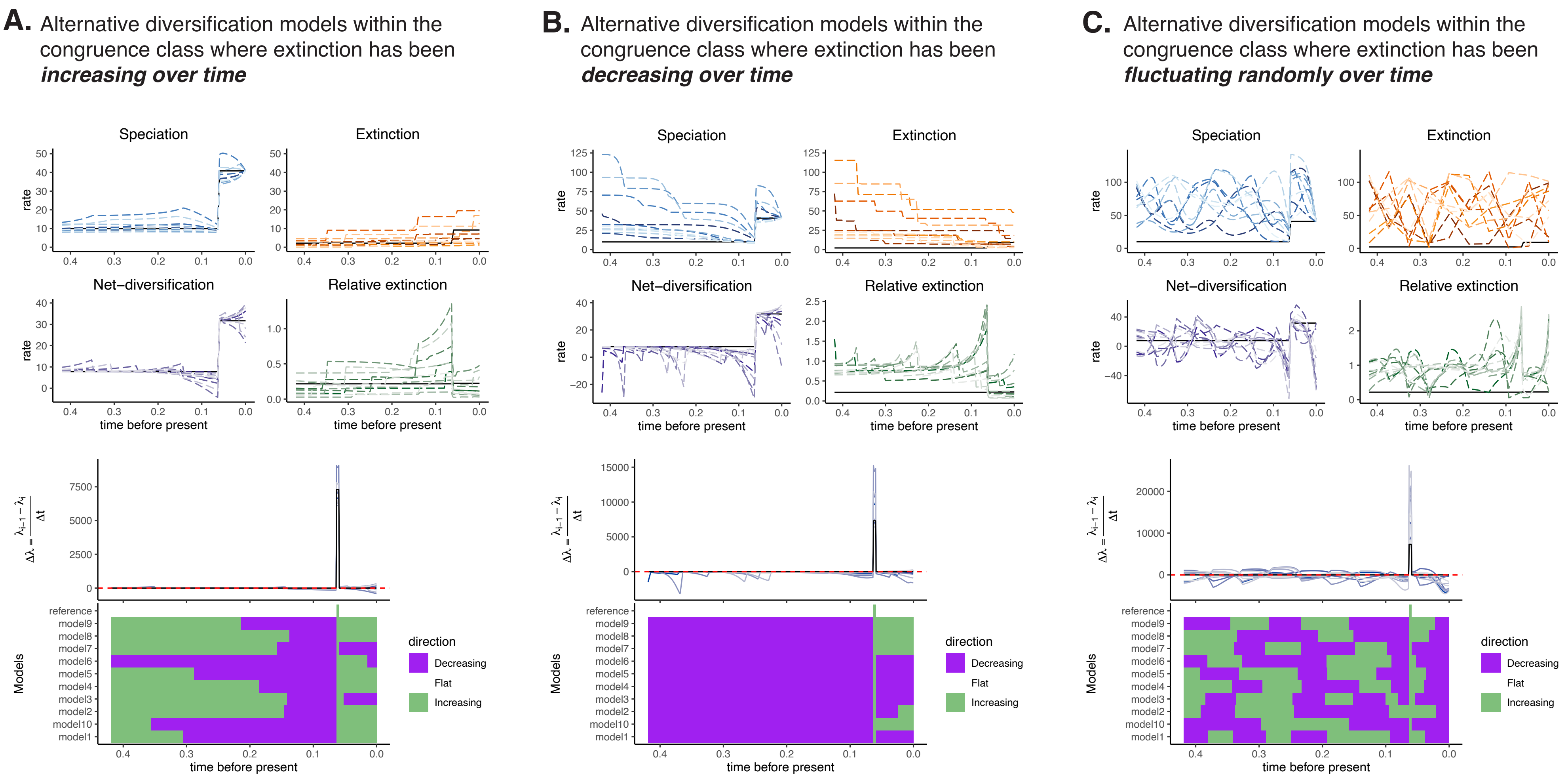

### Supplementary Fig. 9

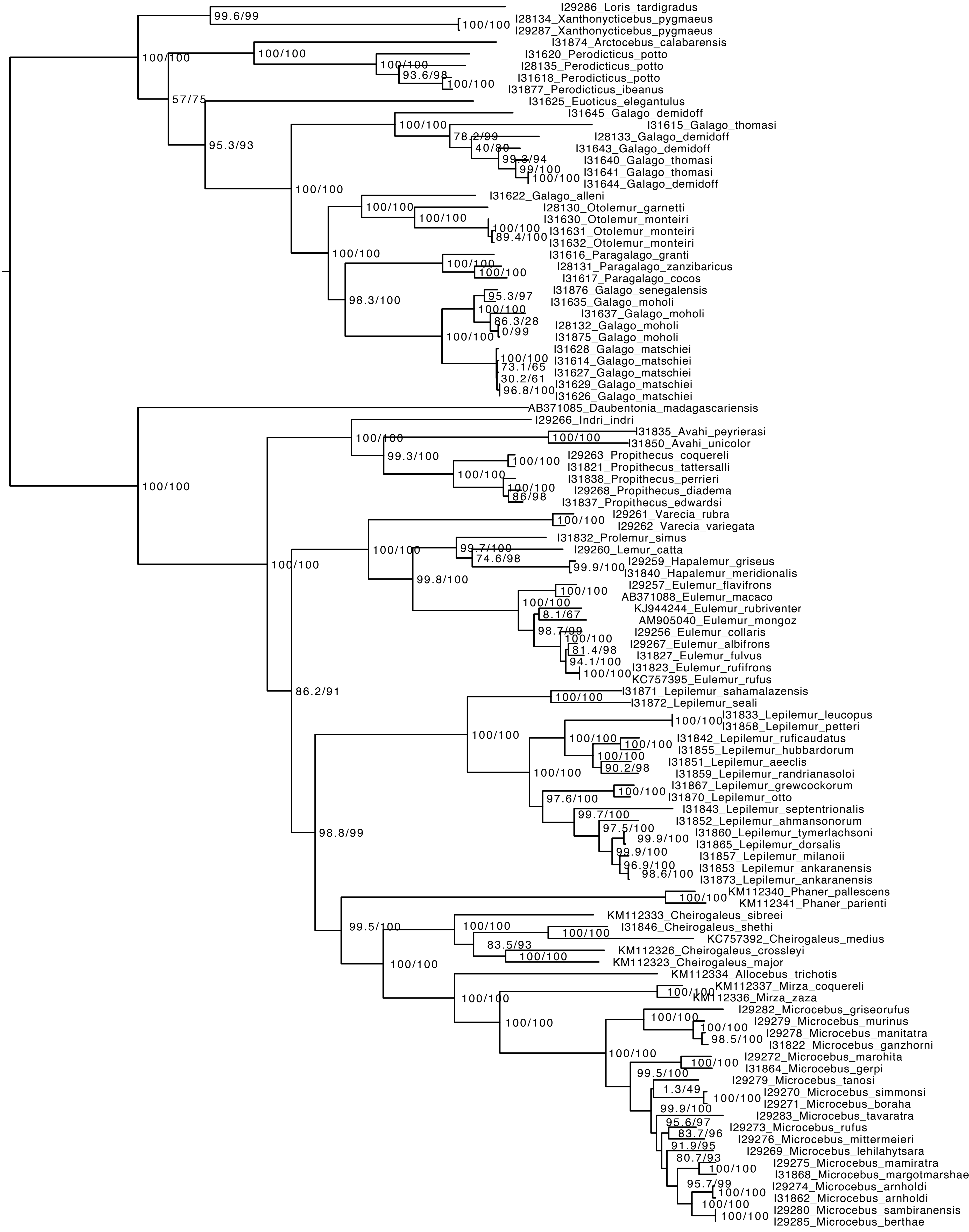

### Supplementary Fig. 10

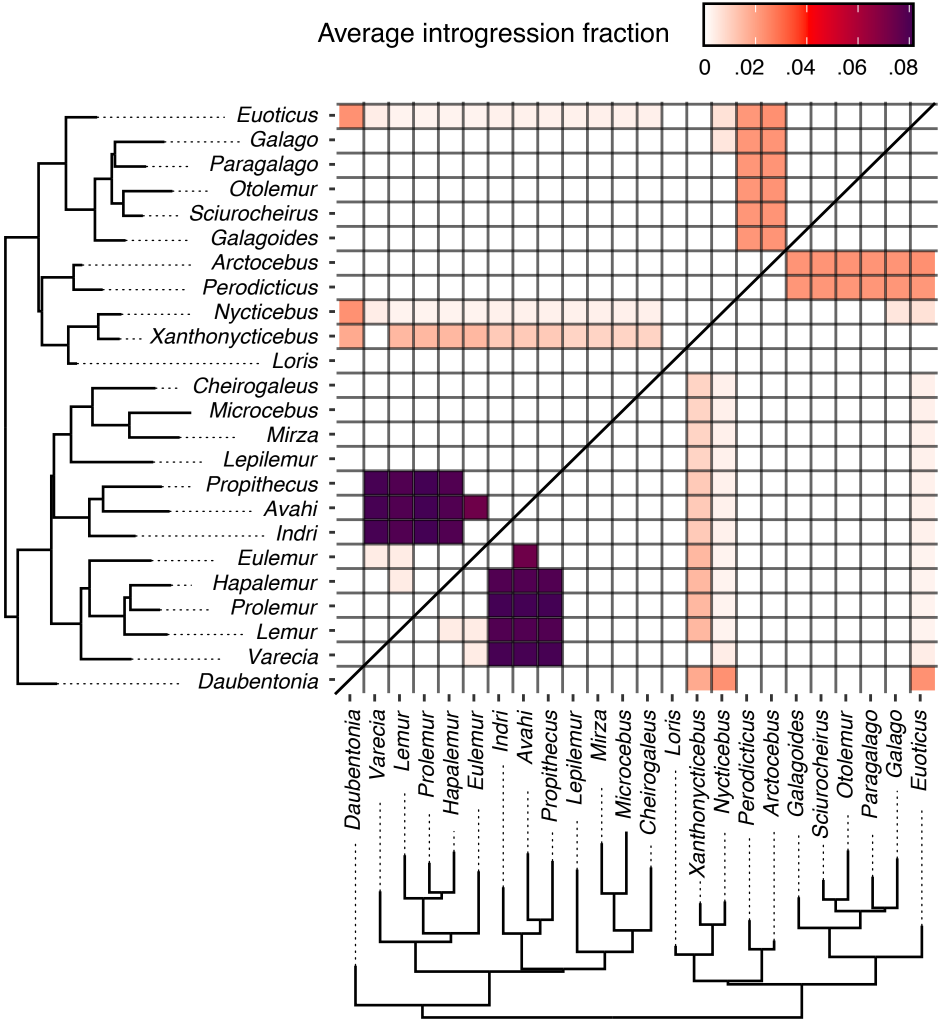

### Supplementary Fig. 11

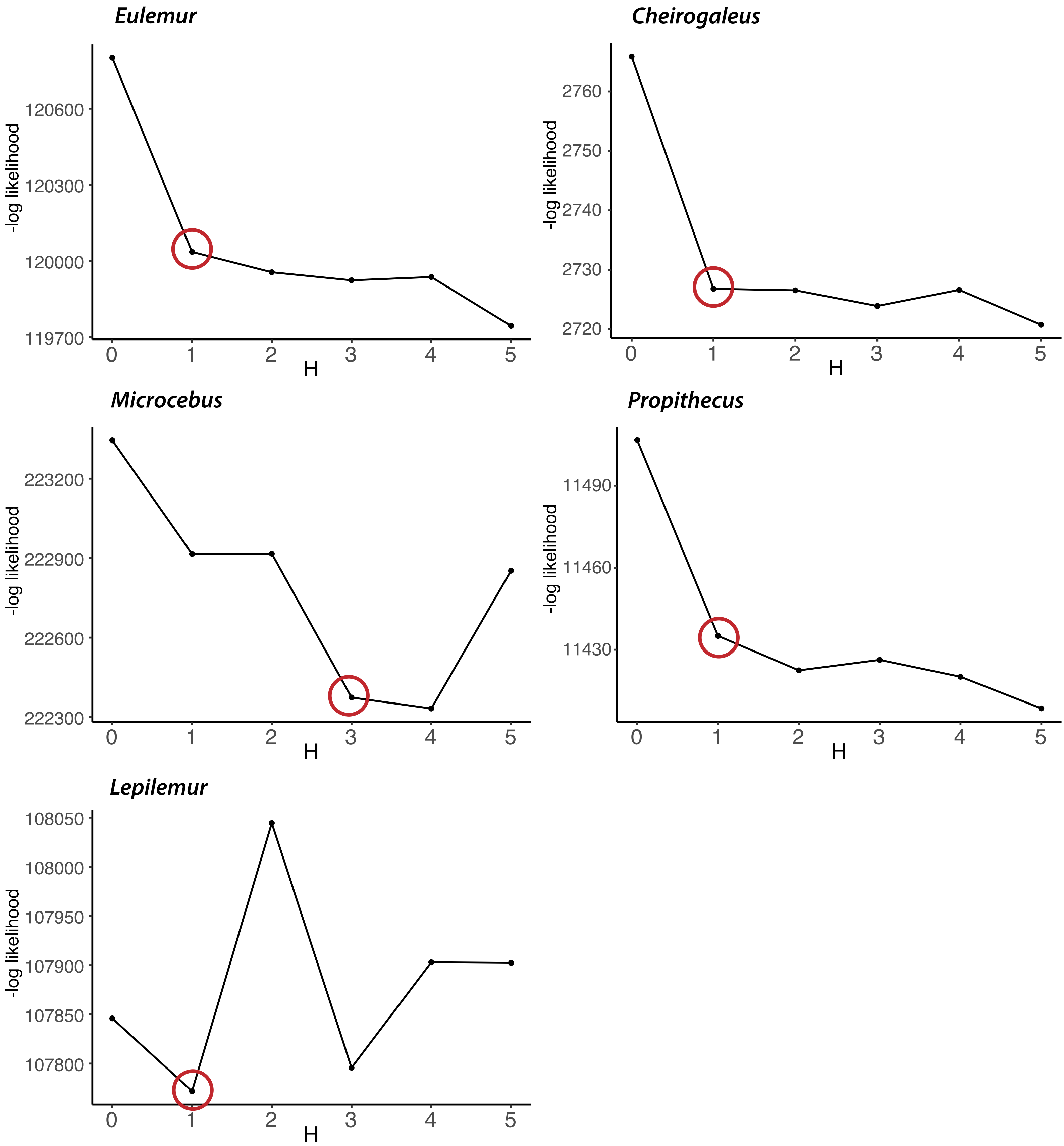

### Supplementary Fig. 12

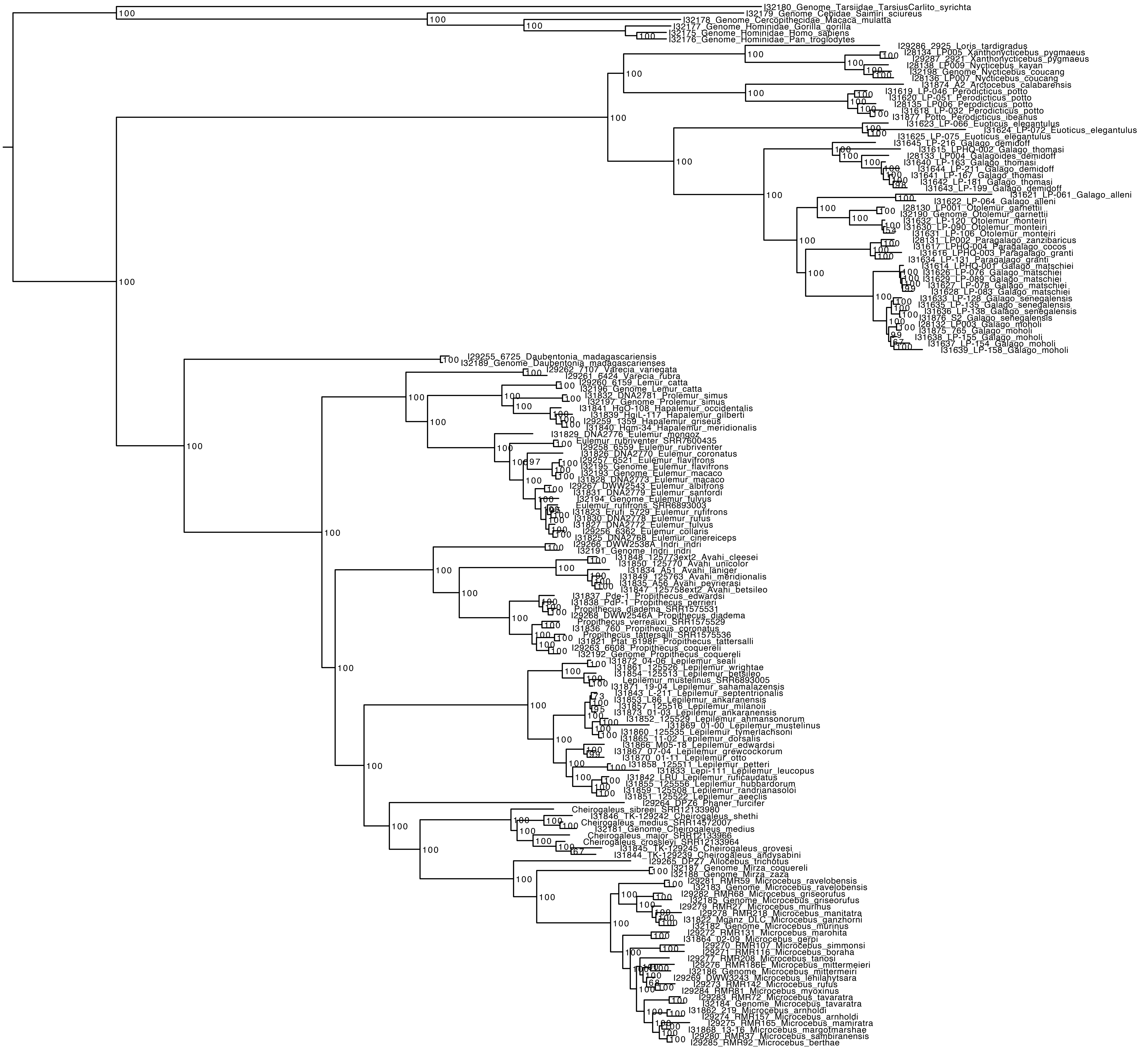

### Supplementary Fig. 13

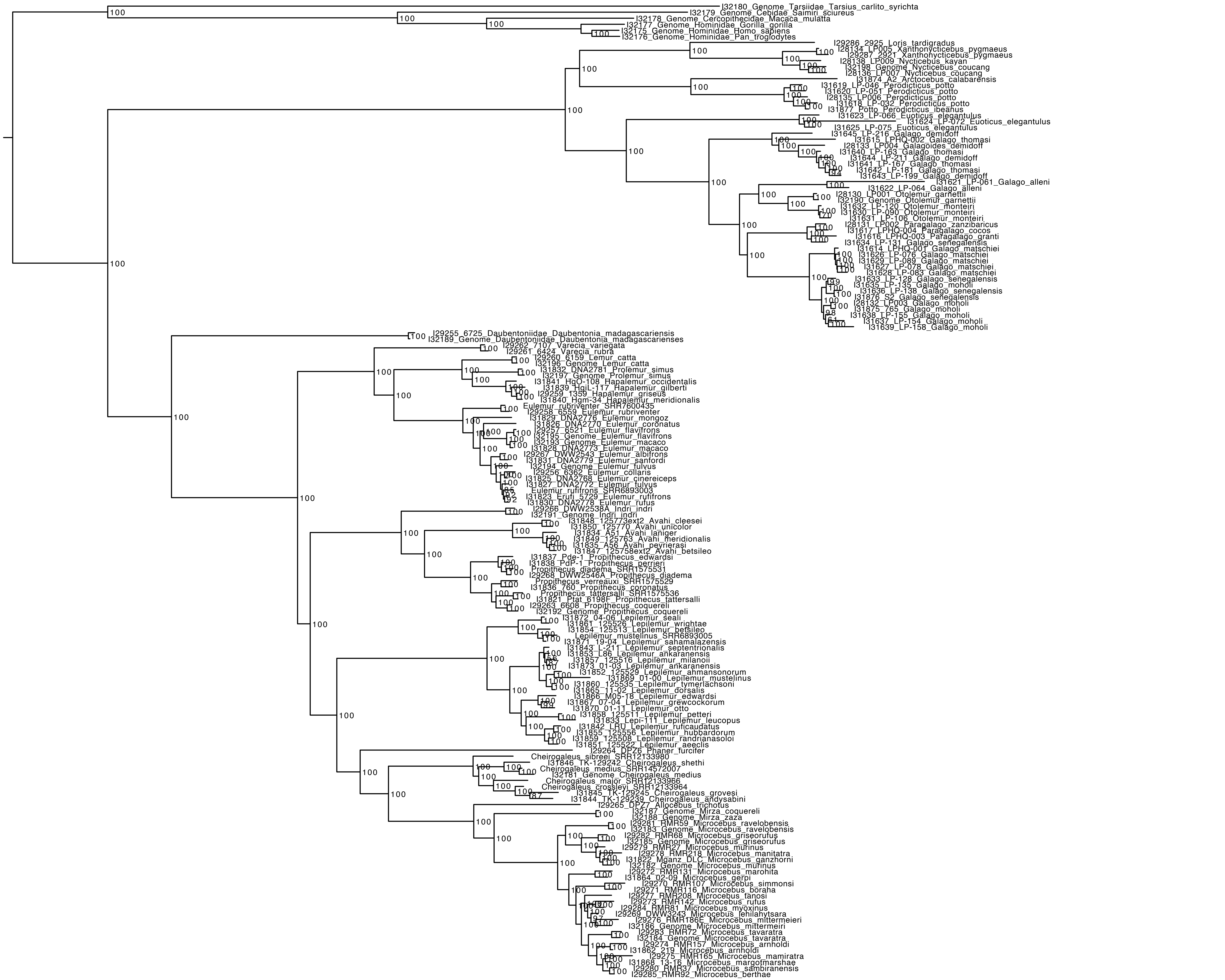

### Supplementary Fig. 14

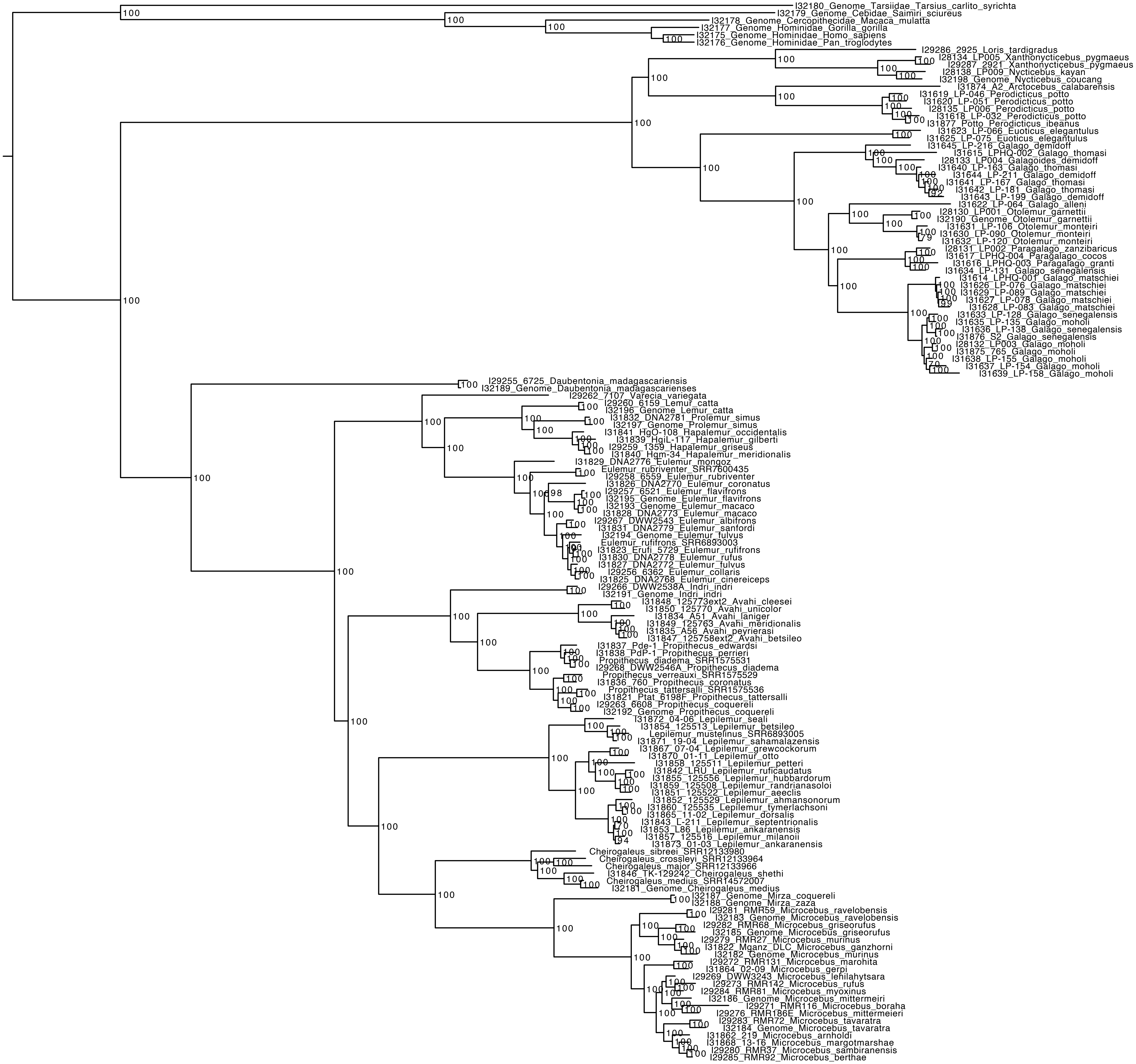

### Supplementary Fig. 15

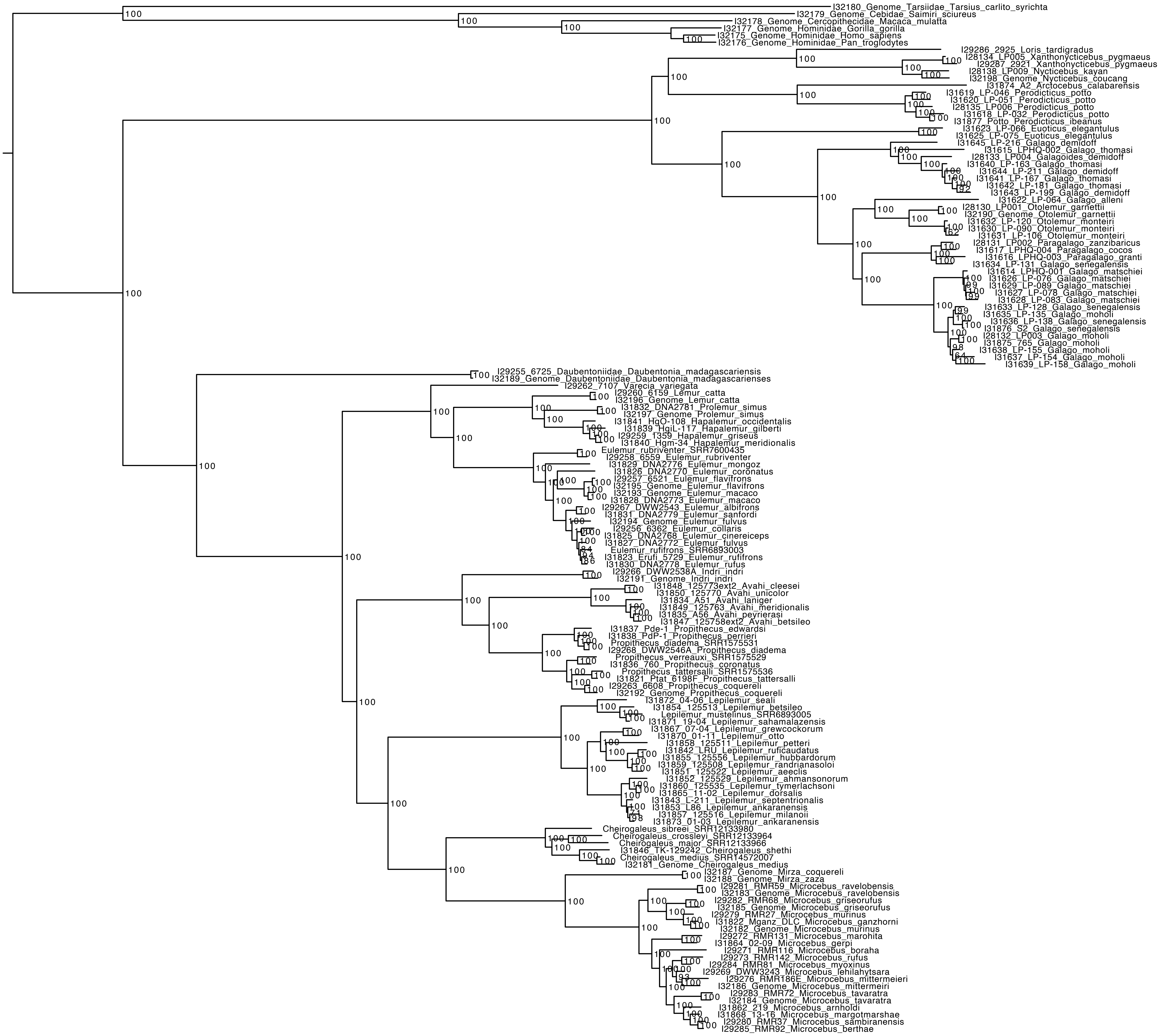

### Supplementary Fig. 16

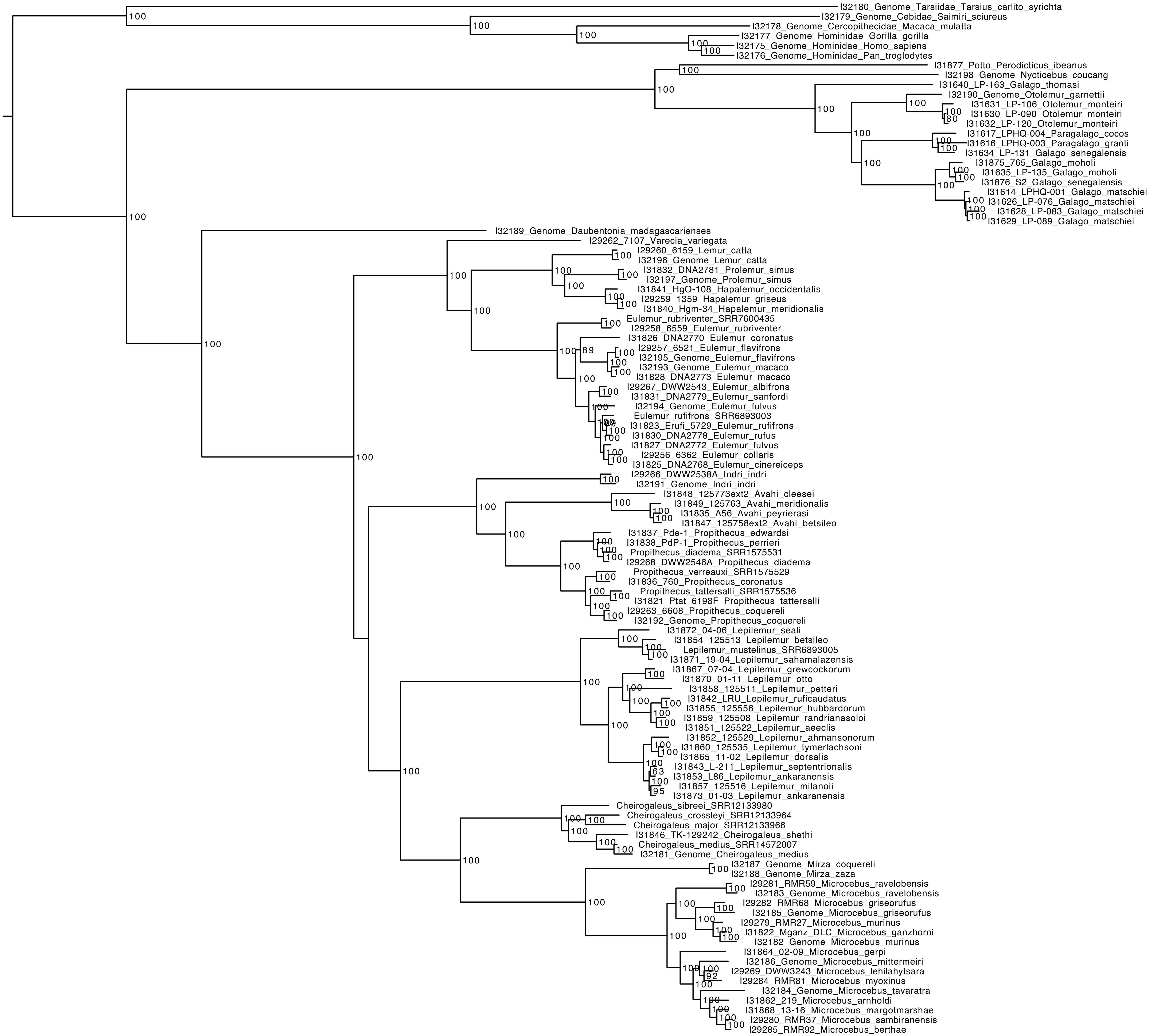

### Supplementary Fig. 17

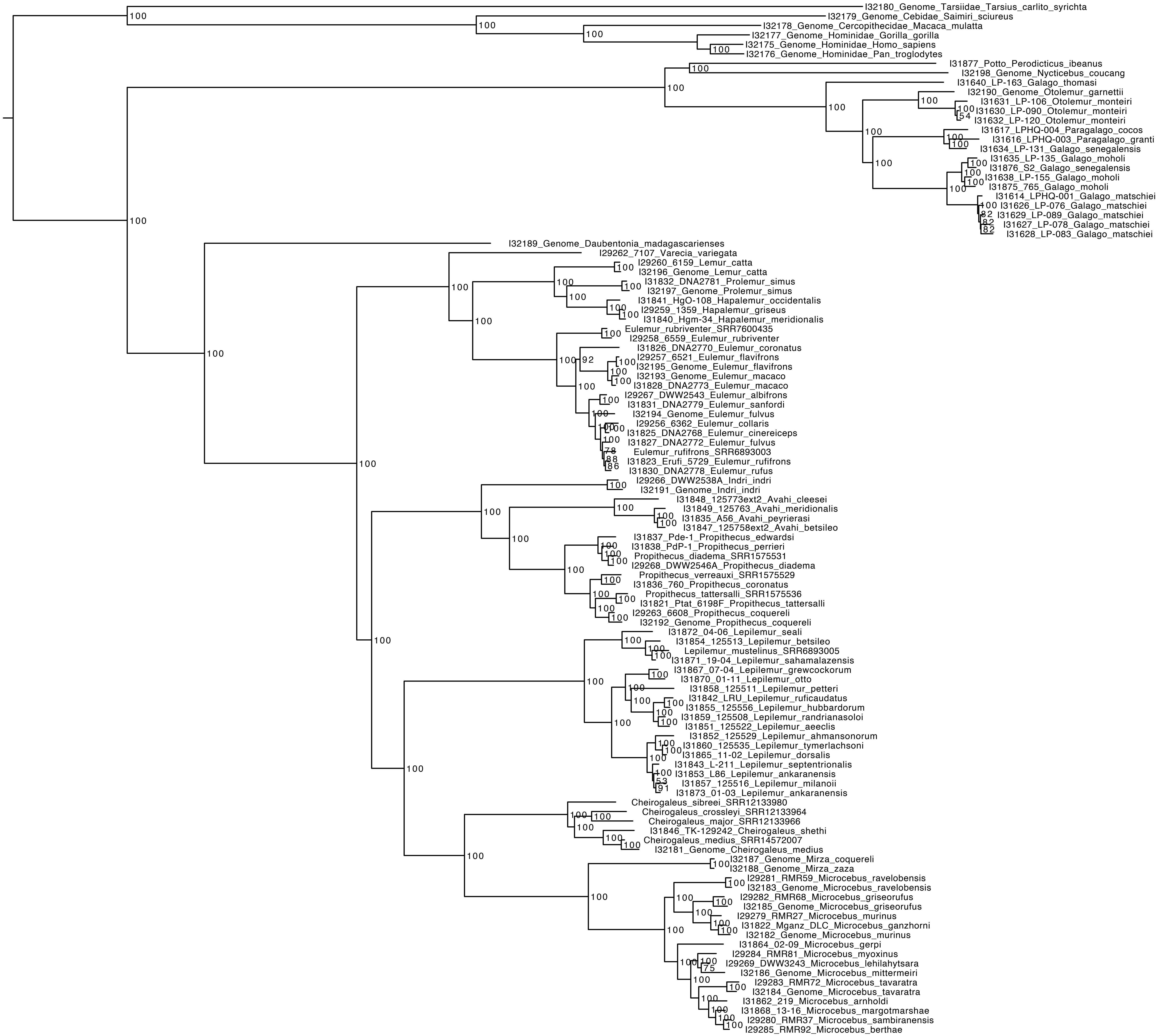

### Supplementary Fig. 18

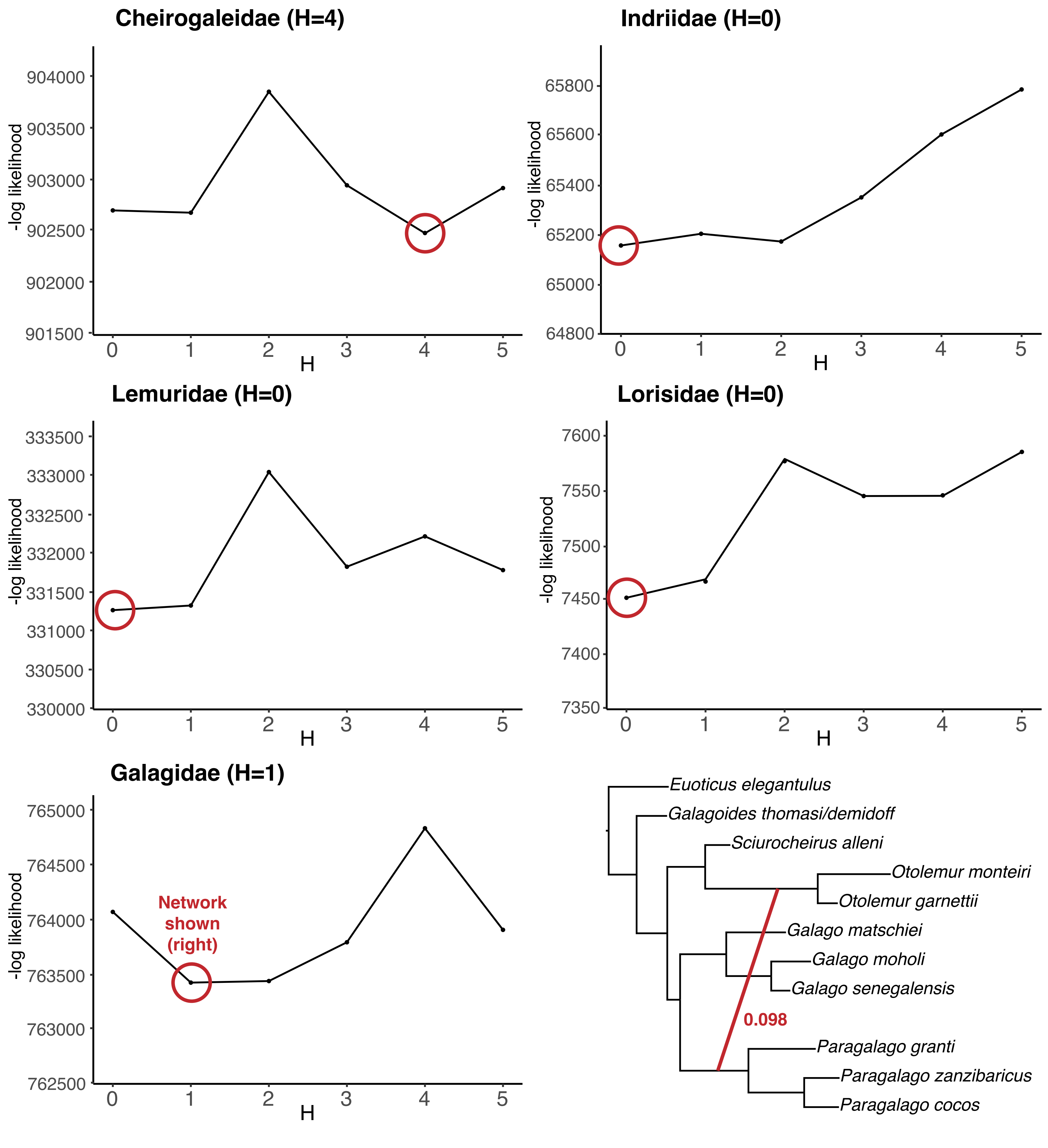
