## Supplementary Fig. 6 for "Multiple bursts of speciation in Madagascar’s endangered lemurs"

A. Combined fossil set with soft root

B. Combined fossil set with hard root

C. Pozzi & Penna fossil set with hard root

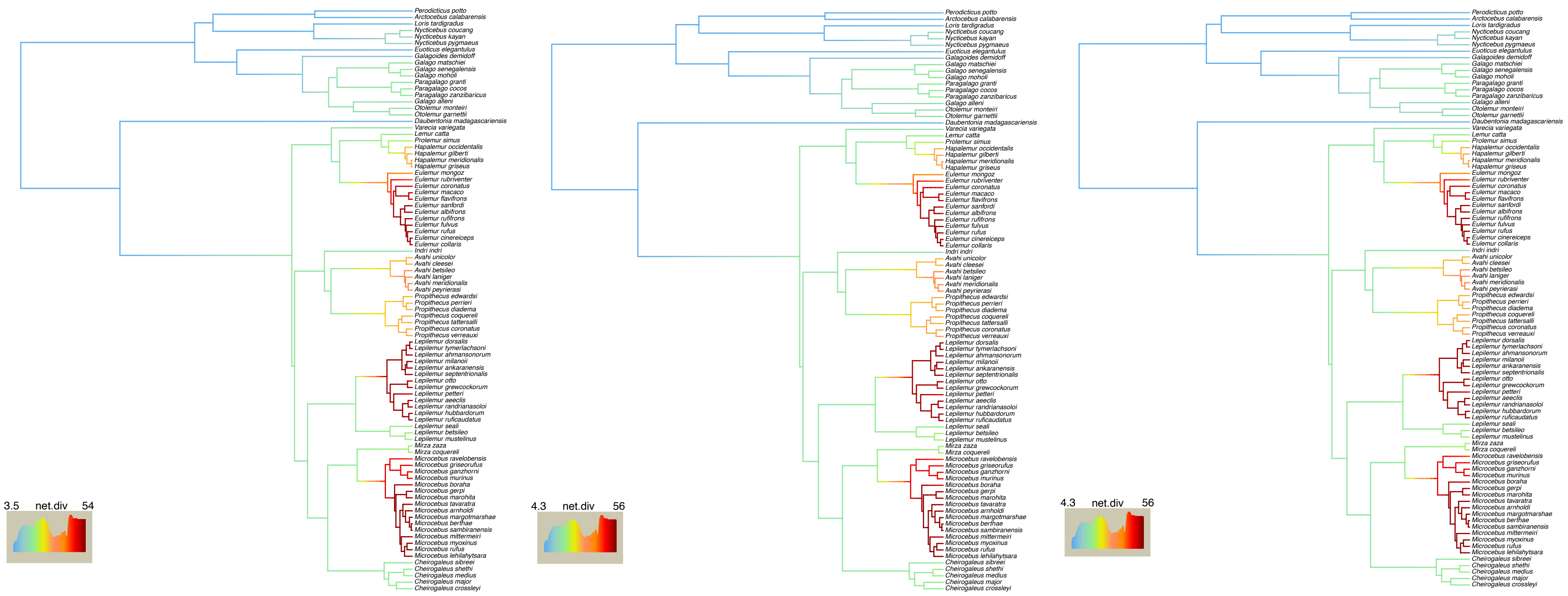
