## Supplementary Methods for "Multiple bursts of speciation in Madagascar’s endangered lemurs"

### Generation of Anchored Hybrid Enrichment data

Samples were sequenced according to the Anchored Hybrid Enrichment (AHE) protocol[1] using a custom probe set (to be released upon publication acceptance). Sequencing and assembly were conducted in two phases: the first phase in 2018 included 42 individuals and the second phase in 2019 included 90 individuals. Both phases were performed on an Illumina NovaSeq 6000 instrument at Florida State University and produced a set of 332 nuclear loci. Quality control and assembly were performed following the same procedure outlined in another recent AHE-based study.[2] We also supplemented our data with previously published whole genome data: six primate outgroups and 28 previously published strepsirrhines (Supplementary Table S7). Briefly, we used the nucleotide Basic Local Alignment Search Tool (BLAST) to identify and pull homologous regions matching our targeted loci from each genome. As a last step, all loci were imported into the software Geneious Prime [3] for a final quality check by eye, with poorly aligned regions being fixed using the Local Realignment tool. Final alignments for each locus were exported from Geneious to nexus, phylip, and fasta files for further analysis.

### Evaluation of the effects of missing data

To evaluate the impact of missing data on phylogenetic analyses, we first created six concatenated fasta files containing:

(1) All loci, all taxa (161 taxa, 1,108,850 bp)
(2) All loci, taxa with >50% missing data removed (144 taxa, 1,108,850 bp)
(3) All loci, taxa with >20% missing data removed (106 taxa, 1,108,850 bp; outgroups with >20% missing data were retained for rooting)
(4) Reduced loci (dropping 37 loci that failed to sequence in lorisiforms and outgroups), all taxa (161 individuals, 969,767 bp)
(5) Reduced loci, taxa with >50% missing data removed (144 individuals, 969,767 bp)
(6) Reduced loci, taxa with >20% missing data removed (109 individuals, 969,767 bp; outgroups with >20% missing data were retained for rooting)

All six datasets were analyzed using IQTree v.2.1.3.[4] Each locus was treated as a separate partition for automatically estimating substitution models, and a maximum-likelihood phylogeny was estimated for each dataset with 1000 ultrafast bootstrap replicates (Supplementary Figs. 11-16).

We observed that missing data had no effect on the overall topology or node bootstrap support values, except that several important genera and species were removed from the datasets with reduced taxa. However, we did observe that many of the taxa with >50% missing data had long terminal branch lengths. Because branch lengths are important in diversification and divergence time analyses, we ran our divergence time analysis (see Materials and Methods) using Dataset 2 (all loci, taxa with >50% missing data removed).
